# Roodmus: A toolkit for benchmarking heterogeneous electron cryo-microscopy reconstructions

**DOI:** 10.1101/2024.04.29.590932

**Authors:** Maarten Joosten, Joel Greer, James Parkhurst, Tom Burnley, Arjen J. Jakobi

## Abstract

Conformational heterogeneity of biological macromolecules is a challenge in single particle averaging (SPA). Current standard practice is to employ classification and filtering methods which may allow a discrete number of conformational states to be reconstructed. However, the conformation space accessible to these molecules is continuous and therefore explored incompletely by a small number of discrete classes. Recently developed heterogeneous reconstruction algorithms (HRAs) to analyse continuous heterogeneity rely on machine learning methods employing low-dimensional latent space representations. The non-linear nature of many of these methods pose challenges to their validation and interpretation, and to identifying functionally relevant conformational trajectories. We believe these methods would benefit from in-depth benchmarking using high quality synthetic data and concomitant ground truth information. Here we present a framework for the simulation and subsequent analysis with respect to ground-truth of cryo-EM micrographs containing particles whose conformational heterogeneity is sourced from molecular dynamics simulations. This synthetic data can then be processed as if it were experimental data allowing aspects of standard SPA workflows, as well as heterogeneous reconstruction methods, to be compared with known ground-truth using available utilities. We will demonstrate the simulation and analysis of several such datasets and present an initial investigation into HRAs.

## 1. Introduction

Cryogenic-sample electron microscopy (cryo-EM) has become an established method to determine three-dimensional (3D) structures of biological macromolecules at resolutions where atomic interpretation becomes possible (Kühlbrandt, 2014; Egelman, 2016). The success of the field has large been attributed to direct electron detectors, improved stability of electron optical components as well as advanced image processing software (Nogales, 2016; Cheng, 2015). One of the remaining challenges in the field is to improve methods for extracting the structural heterogeneity inherent in cryo-EM data to provide meaningful functional insight.

In single-particle averaging (SPA), samples consist of a suspension of biological macromolecules in solution which is then vitrified by flash freezing in liquid ethane. As a result, we presume each particle has a different orientation and conformation. Although the vitrification of the sample is not instantaneous, leaving time for molecules to anneal into a lower-energy state, recent work suggests that a sizeable portion of the conformational states of the protein at room temperature is still populated in the vitrified state (Bock & Grubmüller, 2022).

The final result of a SPA workflow is often a single consensus 3D reconstruction representing the macromolecule of interest.

In such a consensus map, the conformational states of the particle are averaged over, which causes the density, particularly in flexible regions, to be blurred and effectively reduces local resolution. It is therefore standard practice to employ strategies to reduce the heterogeneity in the data by means of discrete 3D classification into structurally homogeneous particle subsets (Scheres, 2016). In cases where the heterogeneity is limited to a discrete set of distinctly different conformations, this strategy can be used to obtain structures of these complexes in their different structural states (Nguyen *et al*., 2016; Zhou *et al*., 2020).

However, subdivision of the cryo-EM data into a discrete number of subsets is not well suited for the description of continuous types of molecular motion, for which an infinite number of subsets would, in principle, be needed. In recent years methods have been developed that aim to more comprehensively explore the conformational heterogeneity present in cryo-EM data. A variety of strategies have been employed to accomplish this goal, which we collectively refer to as heterogeneous reconstruction algorithms (HRAs). One approach, multi-body refinement (Nakane *et al*., 2018), builds on the assumption that largescale conformational changes of a complex can be described by a discrete number of independently moving rigid bodies that themselves are identical across the particle images. After alignment of the individual bodies using signal subtraction and focused refinement, principal component analysis on the relative orientations of all bodies for every experimental image is used to characterise the most dominant motions in the complex. This method is limited by requiring the user to define rigid bodies sizeable enough for refinement within the structure of interest. The estimated conformational heterogeneity is also limited to large domain motion as a result.

Another approach is 3D variability analysis (Punjani & Fleet, 2021) which uses a form of principle component analysis (PCA) to decompose the conformational heterogeneity in a set of particle images into principle modes of motion and a per-particle latent vector. The latent vector is a weighting of all modes of motion present in the particle image. This method offers a way to represent continuous conformational heterogeneity as a continuous manifold, similar to what is discussed by Frank et al. (Frank & Ourmazd, 2016).

Numerous other approaches also aim to embed all 2D projected particle images into a low-dimensional latent space that encodes the heterogeneity in the data by utilising variational autoencoders (VAEs). CryoDRGN (Zhong *et al*., 2021) is an example of such a method, where the encoder and decoder are both implemented as convolutional neural networks and the decoder directly outputs a 3D volume density representation. Whilst for most of these methods the encoder model is similar, there are different choices for the implementation of the decoder model. 3DFlex (Punjani & Fleet, 2023) outputs a deformation field that specifies the way a consensus volume should be deformed to reach a specific state according to the sampled location in the latent space. DynaMight and EMAN2’s e2gmm program opt to model the density as a Gaussian mixture model (Chen & Ludtke, 2021; Schwab *et al*., 2023). The parameters of the mixture model are then the result of passing a latent variable through the decoder.

In many cases these methods are used as a replacement for 3D classification or to filter the particles such that one or more homogeneous datasets remain (Leesch *et al*., 2023; Huang *et al*., 2022; Serna *et al*., 2022; Schoppe *et al*., 2021) and the full potential of HRAs to explore conformational heterogeneity in cryo-EM data is not always utilised. It can be challenging and time-consuming to optimise the model and interpret and validate results obtained with deep learning-based methods. As with any algorithm, the performance of HRAs depends on the hyperparameters used and these may require tuning to achieve the best results. The latent space embedding requires further analysis by clustering or dimensionality-reduction methods to arrive at biologically interpretable results.

We believe HRA development will benefit from thorough benchmarking with known ground-truth data. This will allow the assessment of conformation sampling for both completeness of conformation space and relative populations. It will also allow exploration of the effect of hyperparameter optimisation and aid in interpretation of results.

Simulated data has previously been used in demonstrating various heterogeneous reconstruction methods such as Cryo-DRGN and e2gmm. In these cases a simple linear motion of a structure was simulated. Projection images of the structures were then generated and Gaussian noise was added. Our new approach allows more complex heterogeneity to be modelled by utilising the conformational states sampled from molecular dynamics (MD) simulations. We use Parakeet (Parkhurst *et al*., 2021) software to simulate our datasets. Parakeet employs a multislice approach to propagate an electron wave through a thin amorphous ice layer in which scattering molecules are placed in random orientations. As these micrographs are suitable for processing using conventional 3D reconstruction workflows, it is possible to gain an increased understanding of how the application of each step in the reconstruction workflow, including particle picking and classification, contributes to the final result. We called this simulation and analysis toolkit Roodmus.

In this paper, we first expand upon how the Roodmus toolkit is designed and how it can be utilised to perform the case studies which are subsequently reported. We then demonstrate the effect of including conformational heterogeneity in a synthetic dataset on a conventional SPA reconstruction workflow. We probe the current limitations of the simulation approach through an investigation into how fluence and a radiation damage model affect the consensus reconstructions. Subsequently, we investigate both 3D classification of synthetic data with discrete heterogeneity and explore how conformational heterogeneity is expressed in embeddings of the latent representations provided by two prominent VAE-based approaches CryoDRGN and 3DFlex.

## 2. Results

### 2.1. Roodmus workflow

The goal of the Roodmus toolkit is to generate realistic datasets of synthetic electron micrographs and allow the ground truth information, including particle positions, particle orientations, particle conformations and electron optical parameters to be stored and later mapped to particles in the conventional SPA or HRA workflow metadata files.

The first step (MD sampling, see Fig. 1) serves to acquire a source of structural heterogeneity and to sample atomic coordinate models from it. We suggest sourcing a motion trajectory of a biomolecular complex or other structure of interest. Atomistic MD simulations sampled over a sufficiently long time provide the most detailed observable conformational changes although sampling large domain motion may be too computationally demanding. In these cases various forms of enhanced sampling can be considered as well, such as steered MD, metadynamics or replica-exchange MD (Yang *et al*., 2019). In this work we showcase applications using both atomistic MD trajectories of up to 500 µs, as well as steered MD trajectories that force a large conformational change to occur within time frames of 100 and 200 ps. The left-most panel of Fig. 1 shows an example of a conformational ensemble sampled from a 10 µs MD trajectory of the SARS-CoV-2 spike protein (Shaw, 2020). Large trajectories that are finely sampled can be downsampled to *k* conformations by selecting equidistant timepoints using a Roodmus utility.

**Figure 1.**
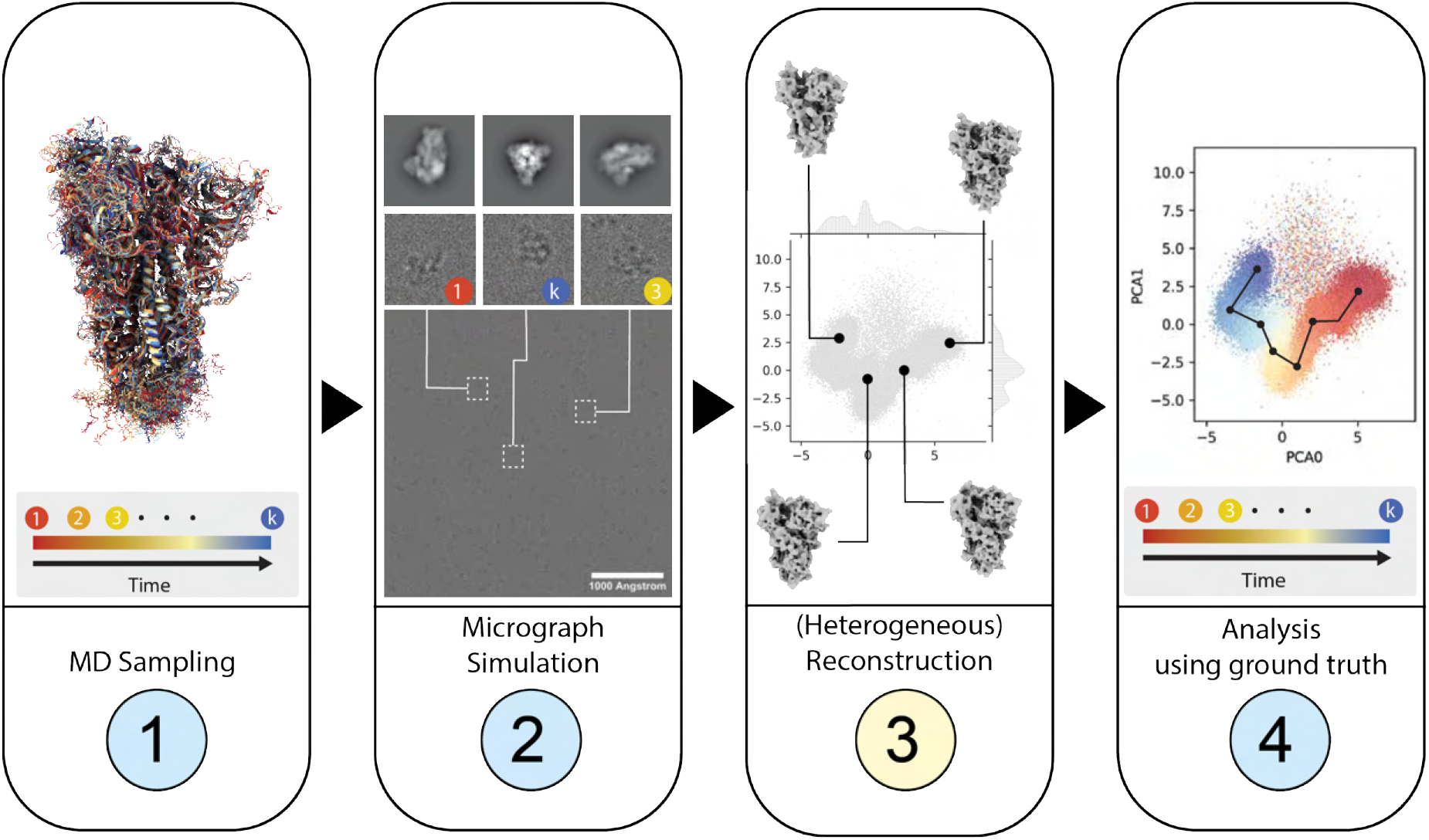
Overview of a 4-step workflow making use of the Roodmus framework. Nodes highlighted in yellow are performed using software external to the Roodmus framework; steps in blue use Roodmus utilities. (1) A user-provided MD simulation is used as the input to the data generation. Timepoints from the trajectory may be sampled via Roodmus, which subsequently simulates micrographs using the Parakeet simulation package in which the conformation of each particle is sampled from these timepoints (2). These synthetic micrographs can be processed using 3D reconstruction software, including HRAs (3). Every step of the resulting reconstruction pipelines can then be analysed using Roodmus utilities to compare reconstruction outputs to ground truth information (4). Roodmus currently supports metadata from CryoSPARC and RELION pipelines as well as CryoDRGN and 3DFlex metadata.

Next, we configure the Parakeet simulation package to create synthetic micrographs. A random selection is made out of the available conformations to build a sample. The positions and orientations of each molecule can be specified at this time or they can be randomly generated by Parakeet, sampling the orientation of each particle uniformly from the SO(3) rotation group. Each particle has a bounding box and cannot overlap with other particles. The sample thickness can also be specified, and can cause particles to overlap in projection when the thickness allows for multiple particles to be placed along the projection axis. Other imaging parameters including fluence, acceleration voltage, defocus and electron optical aberrations can be specified. In our investigations, unless otherwise stated, all datasets created were simulated with a total fluence of 45 *e*^−^/Å ^2^, acceleration voltage of 300 kV, and the Gaussian Random Field ice model (Parkhurst *et al*., 2024) with an ice thickness of 50 nm. In addition, the defocus used in image simulation is sampled from one or more normal distributions, each with an adjustable mean and variance. This is done to reproduce typical experimental defocus distributions resulting from variations in grid planarity and commonly applied focussing protocols during low-dose imaging. Parakeet then propagates the electron wave through the virtual sample and applies the contrast transfer function (CTF) and simulated detector effects to produce a micrograph or dose-fractionated movie.

The synthetic dataset can then be processed using common single-particle averaging workflows or HRAs. Groundtruth information is saved during simulation, including particle positions, orientations and defocus values. This information can subsequently be compared to the metadata output at any stage of processing. Additionally, this information can be compared to the outputs from HRAs. For example, a latent space representation of heterogeneity can be labelled with the time at which that conformation occurs in the MD simulation, as is shown in panel 4 of Fig. 1.

In the next sections we will demonstrate the usefulness of applying synthetic data to all aspects of cryo-EM SPA workflows and show the effects of sample heterogeneity on 3D reconstruction. We will conclude with the application of Cry-oDRGN and 3DFlex to our synthetically generated datasets and compare the results with the known conformational heterogeneity from the simulations.

### 2.2. The effect of increasing heterogeneity in simulated single-particle data

To investigate the impact of heterogeneity on consensus reconstructions, we generated two datasets from a 10 µs MD simulation of the SARS-CoV-2 spike glycoprotein in the partially open conformation (labelled DESRES-ANTON-11021571) (Shaw, 2020). The trajectory consists of 8334 timepoints with a 1.2 ns interval. For the first dataset a single conformation was extracted from this trajectory and used to generate 34 micrographs, containing 10200 particles in total. The second dataset utilised all conformations from the MD trajectory and consisted of 900 synthetic micrographs containing 270000 particles in total. We increased the amount of data generated to ensure that the sampled viewing directions for each conformation are sufficient for later analysis.

Fig. 2A shows the atomic model that was used during simulation of the first dataset. We processed this synthetic data in RELION 4.0 (Kimanius *et al*., 2021) and randomly sampled 8000 of the particles, from which we reconstructed a 2.29 Å resolution map. Except for the lack of motion correction, a standard reconstruction workflow as detailed in section 4.3 was implemented. Local resolution estimation showed little variation except for the expected gradient towards the particle periphery (Fig. 2A), consistent with single conformation and constant atomic displacement factor of the atomic model used for micrograph simulation. The Fourier shell correlation (FSC) curves of the final density map show the expected behaviour with a smooth fall-off to zero. The 3D density appears connected with expected features of sidechains and backbone atoms visible in the structure (Fig. S10). Fig. 2A also shows a plot of inverse resolution versus the logarithm of the number of particles (commonly referred to as a Rosenthal-Henderson or ResLog plot (Rosenthal & Henderson, 2003; Stagg *et al*., 2014)) for this dataset where we obtained a B-factor of 17.8 Å ^2^from the slope of a linear fit. Since the sample is perfectly homogeneous and no B-factor is applied during simulation of these images, this overall B-factor can be solely attributed to errors in processing such as suboptimal alignment.

**Figure 2.**
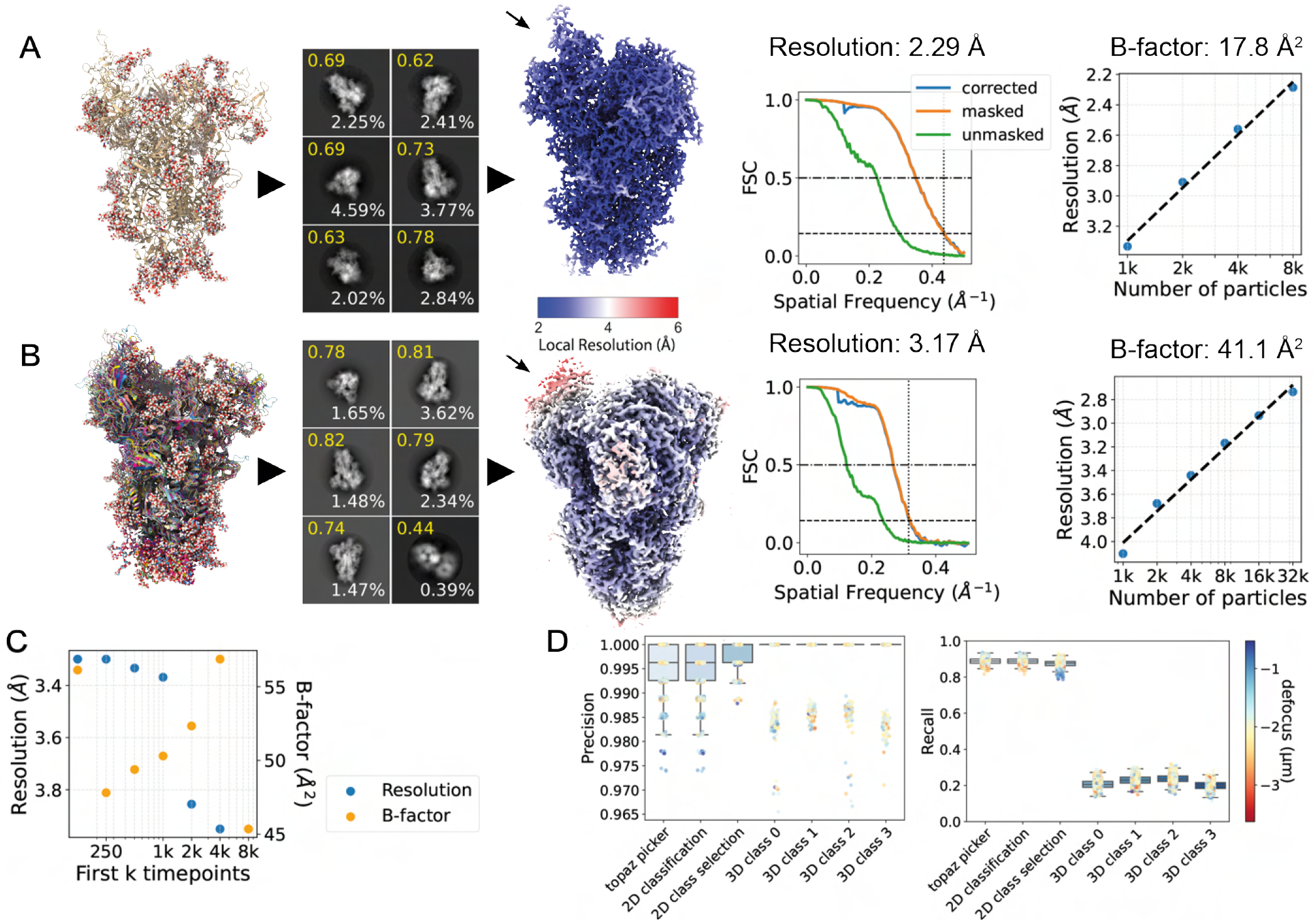
Analysis of reconstructions of single-conformation and conformationally heterogeneous datasets. **A**: Single-conformation dataset. Atomic model (far left), 2D classes (left), 3D reconstruction (middle), FSC curve (right) and ResLog analysis (far right). For 2D classes, the ‘class score’ from automatic class ranking in RELION is shown in yellow and percentage of particles in the class in white text. Density map is coloured by local resolution. The arrow indicates the RBD. **B**: Conformationally heterogeneous dataset. Order of panels as in (**A**); the atomic models shows an ensemble of structures from conformational sampling. **C**: Resolution (blue, left side axis) and global B-factor (orange, right side axis) versus number of included conformations in the dataset. The number of particles is constant for each dataset. **D**: Precision and recall for various steps of the image processing workflow for the conformationally heterogeneous dataset.

The second dataset of synthetic micrographs were then preprocessed (Fig. S9) using the same workflow as the single-conformation dataset. In order to compare a consensus reconstruction of this dataset with a consensus reconstruction of the previous single-conformation dataset we similarly selected 8000 particles randomly before refinement. The resulting density map is shown in Fig. 2B. We now observe a lower global resolution of 3.17 Å as well as larger differences in local resolution compared to the single conformation dataset. In the single-conformation dataset we measure a mean local resolution of 2.2 Å with a variance of 0.61 Å ^2^, while the conformationally heterogeneous dataset has mean 3.0 Å with variance 1.1 Å ^2^. The largest difference in resolution is observed in the flexible remote binding domain (RBD), which is in a partially open conformation in this MD trajectory. Performing a ResLog analysis shows the overall B-factor has increased to 41.1 Å ^2^.

We further investigated the relation between heterogeneity present in the dataset and the global resolution of a reconstructed density map. To this end we created 7 subsets of 4000 particles from the multiple conformation dataset. Each subset only includes particles from the first *k* timepoints of the MD trajectory with *k ∈* (125, 250, 500, 1000, 2000, 4000, 8000). Fig.2C shows the global resolution and 3D refinement-estimated B-factor as a function of *k*. As expected, the global resolution decreases and the B-factor increases as the conformational heterogeneity contained in a subset increases. The overall resolution was lower compared to the consensus reconstructions of the single and multiple conformation datasets (possibly caused by omission of CTF refinement). The B-factor data also shows unexplained outliers which might result from the low number of particles used in refinement and stochastic nature of refinement.

As ground truth particle positions, orientations and CTF parameters are known, we next investigated how well the corresponding estimates from the 3D reconstruction workflow agreed with the ground truth. Fig. S1A plots the estimated defocus values against the ground truth defocus for each micrograph, from which we computed a correlation coefficient of 1.0 to 6 significant digits. We also compared ground truth particle orientations to estimated orientations during 3D refinement with distributions of elevation and azimuthal angles shown in Fig. S1B and C. The ground truth distribution of viewing directions is uniform, whereas the estimated orientations show a peak which may indicate misalignment for some particles.

To analyse particle picking we matched each picked particle with the closest truth particle within a radius of 50 Å . We define true positive picks as those picked particles which were successfully matched to a truth particle and false positives as those which were not. From these definitions we can compute particle picking precision and recall for a set of picked and ground truth particles. Precision and recall for various steps of processing of the heterogeneous dataset are plotted in Fig. 2D on a per-micrograph basis. Particle picking is nearly perfect; a trained Topaz (Bepler *et al*., 2019) model can pick 88.8% of all particles with 99.5% precision. As we will discuss in section 2.4, we also simulated a steered MD trajectory of only a monomer of the SARS-CoV-2 spike protein. Compared to the trimer, particle picking was found to be more difficult for this smaller structure. Fig. S1E shows particle picking precision and recall for the smaller structure and here Topaz finds 92.3% of all particles with a precision of 82.6%. The plot also shows that, in general, the recall is higher in micrographs with a larger defocus, while the precision is lower for both the blob picker and subsequent Topaz picking.

### 2.3. Radiation damage and fluence

One problem with the data simulation is the lack of frequency dependent attenuation of the signal, which would be expected for experimental data as a result of beam-induced motion, radiation damage and detector response. Parakeet offers the option to model the detector response using a detective quantum efficiency (DQE) model which attenuates amplitudes in a frequency-dependent manner according to tabulated values based on a Falcon4 detector at 300 keV. The DQE also depends on the electron flux density; in our still image simulation, we used 5 *e*^−^/Å ^2^/s. In addition, to account for the effect of radiation damage on the Coulomb potential of the sample, a beam damage model is implemented as a Fourier filter, which effectively convolves the electrostatic potential with a Gaussian function whose variance is proportional to a B-factor. This B-factor *B* = 8*π*^2^*D*_*E*_ *S*_*E*_ depends linearly on the fluence *D*_*E*_ and a sample-dependent sensitivity coefficient *S*_*E*_ with unit Å ^4^/*e*^−^ (Parkhurst *et al*., 2021). In our application, the B-factor model serves to progressively blur the atomic potential for each subsequent frame during simulation of a dose-fractionated movie in order to recapitulate empirical observations of progressive beam damage in which the first few frames of a dose-fractionated movie are least affected by radiation damage and the later frames progressively lose high-frequency information (Grant & Grigorieff, 2015).

To illustrate the effect of the radiation damage model, in Fig. 3A we compare exemplary power spectra from two simulated datasets. The first is from the heterogeneous SARS-CoV-2 trimer dataset introduced in section 2.2 which contains non-fractionated micrographs without radiation damage (-RD). The second is computed from the sum of a 30-frame dose-fractionated movie with radiation damage enabled (+RD). In both cases, the DQE model was enabled and the total fluence was 45 *e*^−^/Å ^2^. Micrographs corresponding to the power spectra are shown in Fig. S2A and S2B. To comparatively evaluate the effect of the radiation damage model on the 3D reconstruction, we once again computed a consensus reconstruction using 8000 randomly selected particles from the SARS-CoV-2 spike glycoprotein simulated with radiation damage, yielding a reconstruction with a global resolution of 3.6 Å as shown in Fig. S2C.The FSC curve is plotted along with the corresponding FSC curve of an analogous reconstruction without radiation damage model (see Fig. 2B). ResLog analysis, shown in Fig. 3C results in a B-factor of 54.1 Å^2^, compared to 45.9 Å^2^without radiation damage, suggesting that radiation damage has a relatively minor effect under the conditions used in our simulations. We note that no beam-induced motion model is currently applied in either of the datasets.

**Figure 3.**
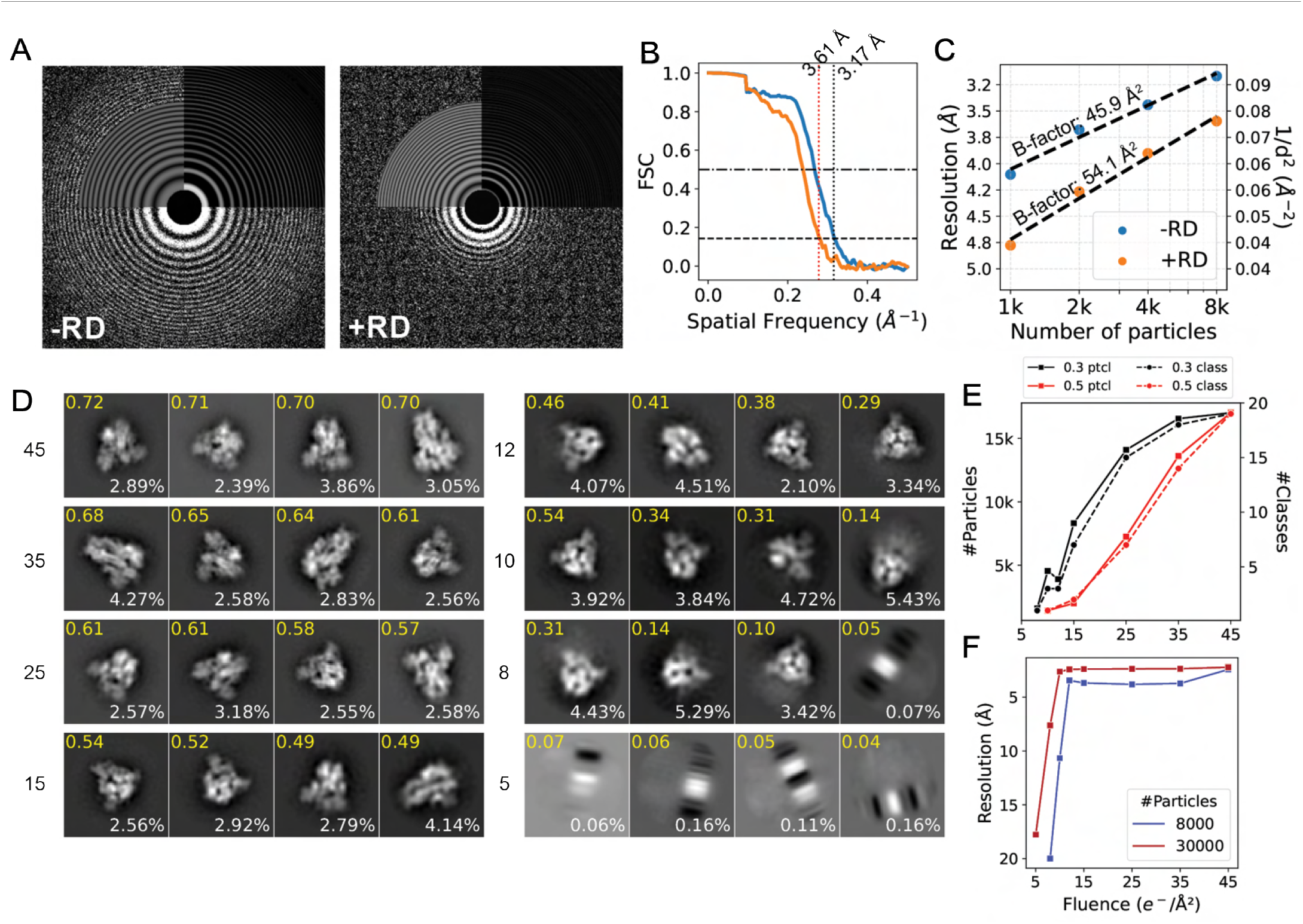
Radiation damage and fluence. **A**: Power spectra of two micrographs simulated without (left) and with radiation damage (RD; right). **B**: FSC curves of reconstructions obtained with 8000 particles from datasets without (blue) and with (orange) radiation damage. **C**: ResLog analysis of datasets simulated without (blue) and with (orange) radiation damage; B-factor from linear regression are shown. **D**: 2D class averages from eight datasets with fluence ranging between 5 and 45 *e*^−^/ ^2^. The four best classes are shown along with their *relion class ranker* score in yellow and the percentage of particles in each class in white. **E**: The number of particles (left axis, solid line) and 2D classes which pass the threshold (right axis, dashed line) of 0.3 (black) or 0.5 (red) for the predicted class scores from RELION. **F**: Resolution vs. fluence plots using either 30000 (red) or 8000 (blue) particles.

In addition to radiation damage, we also investigated the effect of fluence on various steps of the reconstruction process. We simulated a series of eight datasets with fluences of (45, 35, 25, 15, 12, 10, 8, 5) *e*^−^/Å^2^, each consisting of 100 micrographs containing 30000 particles of the SARS-CoV-2 spike protein in total.

We picked particles from the micrographs in each dataset using the Laplacian of Gaussian (LoG) picker in RELION, followed by 2D classification with the *relion class ranker* (Kimanius *et al*., 2021). We found that the quality score of the classes decreased with lower fluence. Fig. 3D plots the 4 highest scoring classes for each dataset. Visually, classes obtained from datasets with lower fluence also appear more blurred, or poorly centred. Fig. 3E plots the number of 2D classes and constituent particles retained after 2D class selection using the automated class ranker with threshold values of 0.3 or 0.5. The plot indicates that for low SNR data the main bottleneck in the reconstruction of these simulated images becomes the alignment and 2D classification. To further illustrate this, we used the groundtruth particle positions to compute the average distance between each picked particle and its nearest ground-truth particle location. The distribution of these distances is plotted in Fig. S2F, which shows that the accuracy of the picked particle locations decreases with decreasing fluence. This may contribute to the 2D classification failing to centre the classes for low fluence datasets.

We obtained reconstructions of each dataset using groundtruth coordinates of the particles and the same reference density. The resolution of these density maps is plotted in Fig. 3F as a function of fluence. We found that the resolution remains roughly constant until the fluence drops below 10 *e*^−^/Å^2^. In the absence of radiation damage the high-frequency information in the images is over-represented compared with experimental data, causing the resolution to remain constant until the alignment fails. In all subsequent case studies, we opted to use 45 *e*^−^/Å^2^.

### 2.4. Discrete classification of simulated dataset with multiple conformations

Next, we applied our pipeline to analyse 3D classification of a synthetic dataset comprised of both discrete and continuous heterogeneity. For this purpose we created a dataset with two discrete structural states by mixing conformations from the previously used DESRES-ANTON-11021571 trajectory, in which the spike glycoprotein is in a partially open conformation (single RBD up), with conformations from the DESRES-ANTON- 11021566 trajectory (Shaw, 2020), in which the protein is in the closed conformation (all RBDs down). We simulated 400 micrographs containing 60081 particles from the open state trajectory and 59919 particles from the closed state trajectory and processed them to obtain an initial consensus density map. From there we performed 3D classification and analysed the distribution of particles in the open and closed conformations in each 3D class. Fig. 4A shows this distribution for the case of two classes (I), three classes (II) and ten classes (III). We observed that using two classes did not result in a clean split between the trajectories, as 78.2% of particles in class two originated from the closed state conformation and 21.8% from the open state compared to 2.1% closed state and 97.9% open state for class 1. In the case of three classes, classes 1 and 3 neatly distinguished between the open and closed states respectively, but class 2 was a mix of both the closed and open states (62.2% closed state, 37.8% open state). In the case of ten classes we found that all classes contained mostly particles from one of the trajectories, with class five being the least uniform (88.3% of particles originated from the closed state).

**Figure 4.**
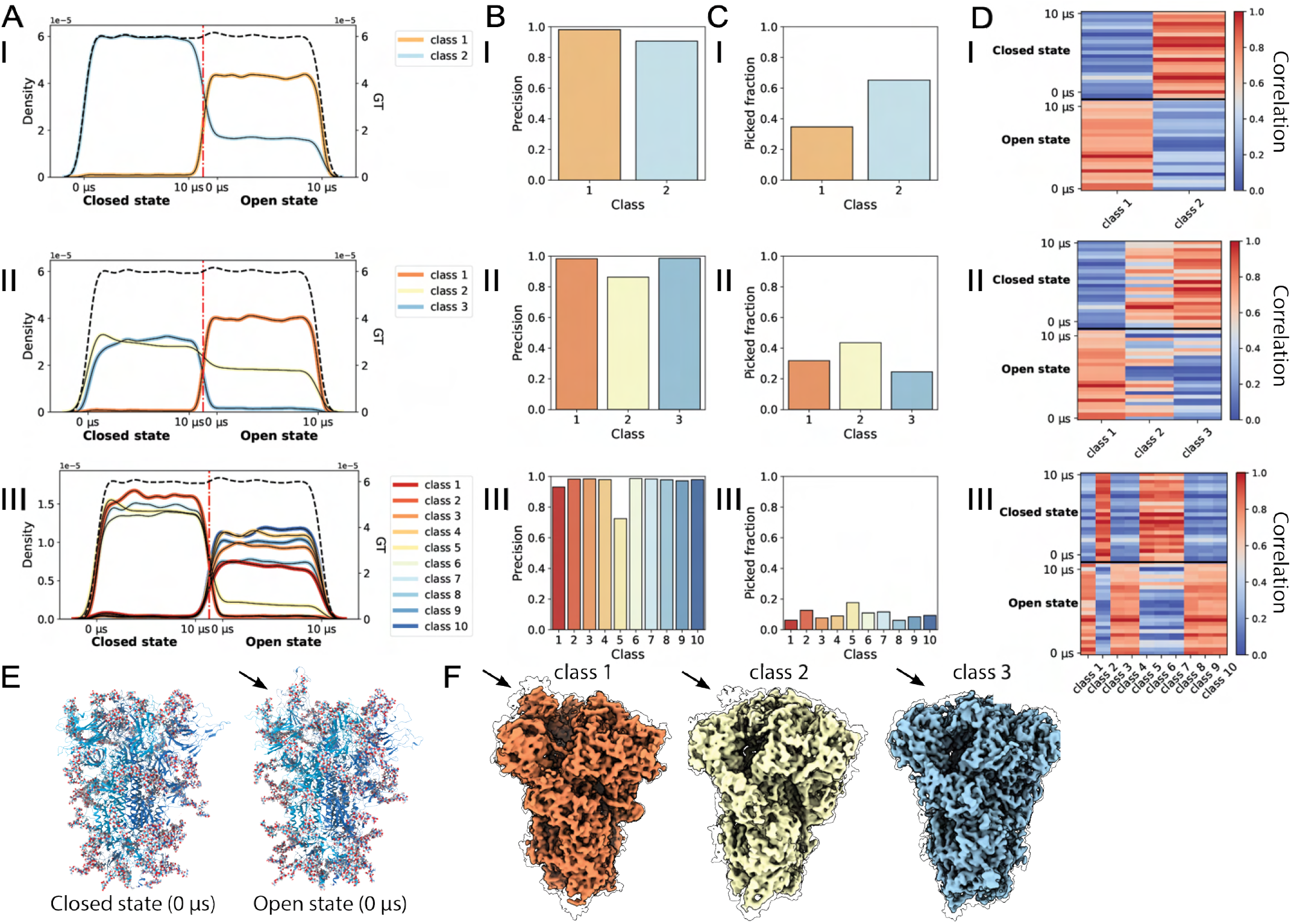
Discrete classification of conformationally heterogeneous synthetic data. **A**: The distribution of particles in each class over the 10 µs trajectories of the spike protein in closed and open states in the case of 2 classes (I), 3 classes (II) and 10 classes (III). **B**: The precision of each 3D class. **C**: The fraction of particles in each class. **D**: A correlation heatmap where each class (x-axis) is correlated against snapshots taken from the MD trajectories of the closed and open states (y-axis). The value plotted is the real-space correlation between class and MD timepoint, normalised per column. **E**: A single conformation from the closed and open states of the protein. **F**: The 3 classes resulting from classification II; transparent contours outline confidence maps thresholded at 5% FDR. The arrow highlights the RBD.

We found that false positive (FP) particles tend to accumulate in one or two classes during classification, which is also typically the class with the most particles and the most equal contribution from both trajectories. This is illustrated by Fig. 4B which shows the precision of the particle set for each 3D class whilst Fig. 4C shows the distribution of the total number of particles over the classes. From refinement of each class of the 3-class classification example, we found the resolutions to be 2.54, 2.78 and 2.6 Å respectively. Class 2, despite containing more particles, resulted in a lower resolution density map.

We then measured the real-space map-to-model correlation using the density map reconstructed from each class after refinement and the backbone atoms of atomic models from 50 evenly spaced timepoints from the MD trajectories of the closed and open states. Fig. 4D shows a heatmap of the normalised correlation between each pair of atomic models and density maps. When using two classes, we found that each class has the highest correlation with atomic models from the trajectory comprising the majority of its constituent particles. When using three classes, we found that this was also true for class 1 (98.5% of particles originated from the open state and the density map correlates more strongly with atomic models of the open state), and for class 3 (95.1% of particles originated from the closed state and the density map correlates slightly better on average with atomic models in the closed state). Class 2, containing particles from both open and closed state, correlates strongly with atomic models in the closed state. In the case of ten classes, we again find that each class correlates strongly with atomic models from the trajectory comprising the majority of its constituent particles. Unnormalised versions of these correlation heatmaps (Fig. S3A) show in addition that there is a large difference between classes. Visual inspection of the density maps for these 3 classes, shown in Fig. 4F shows that class 1 resembles the open state as evidenced by the RBD (see arrow) being in the up conformation. While class 2 and class 3 do not show clear evidence for the RBD in the up conformation even at low density thresholds, false discovery rate (FDR)-controlled confidence maps (Beckers *et al*., 2019) computed for all reconstructions reveal detectable signal for the RBD in the up conformation in class 1 and 2 but not class 3, consistent with the class distributions determined in 4A.

In addition to the synthetic dataset created by mixing the 10 µs trajectories of the molecule in the closed and partially open state we performed a short steered MD simulation interpolating between the closed and partially open state. To reduce computational complexity we only selected one monomer of the complex out of PDB models 6xm4 (open state, RBD up) and 6xm5 (closed state, RBD down) (Zhou *et al*., 2020). We then used harmonic restraints to drive the molecule from open conformation to closed using openMM (Eastman *et al*., 2017). Out of the resulting 100 ps trajectory we sampled 10000 conformations, which were used to create 800 micrographs comprising a total of 200000 particles. As discussed in section 2.2, the smaller size of the particles meant preprocessing was more difficult and more particles were needed. Ab-initio model building with 4 classes was done in CryoSPARC using 213486 picked particles, of which 168556 were true positives. Similarly to the SARS-CoV-2 spike glycoprotein trimer, we again observed 3D classes distinguishing between the open and closed conformation of the molecule. The distribution of the particles in each class over the steered MD trajectory is shown in supplementary Fig. S3B. Classes 2 and 3 had substantially lower precision (0.61 and 0.62 compared to 0.99 and 0.97 for classes 0 and 1) and failed to produce initial models that resembled the spike monomer. These classes also contained particles from the entire trajectory, in contrast to classes 0 and 1 which contained mostly particles from the last 60 ps and the first 40 ps of the trajectory respectively. All 3D classes, refined density maps and their comparison to atomic models are shown in supplementary Figs. S3D to S3F.

### 2.5. Continuous heterogeneity in the latent space is well-preserved

Synthetic data simulated with Roodmus is by design well-suited for heterogeneous reconstruction methods that aim to derive a latent representation of the images. In this section we showcase the application of two such methods - CryoDRGN (Zhong *et al*., 2021) and 3DFlex (Punjani & Fleet, 2023). We apply the former to a synthetic dataset of the SARS-CoV-2 spike trimer glycoprotein in its partially open conformation as well as the SARS-CoV-2 replication transcription complex (RTC) and the latter to steered MD simulations of the SARS-CoV-2 spike glycoprotein (monomer) and the protein Complement C3, a component of the human complement system. These datasets differ in molecular mass, dimensions and number of the particles, the magnitude of the conformational change, the timescale simulated and the coarseness with which states are sampled.

The open-state SARS-CoV-2 dataset is the same as the conformationally heterogeneous dataset used for the consensus reconstructions in section 2.2. The final cleaned data consists of 236079 particles taken after performing a 3D refinement in RELION. The default 8-dimensional latent space was used for CryoDRGN training. In Fig. 5A we show a 2D embedding of the latent space using PCA for dimensionality reduction. FP particle picks, which make up 0.03% of the dataset are coloured red and are spread throughout the latent space. Fig. S4A shows a density plot that visualises how the latent space embeddings do not form distinct clusters. We then use ground truth information to colour each particle according to the timepoint of the MD trajectory from which it originated as shown in Fig. 5B. Conformations that are close together in time were also found to cluster together in the latent space embedding. We grouped the MD trajectory into 50 batches of conformations, each representing a contiguous 2% of the total number of timepoints sampled in the ground truth MD trajectory. By calculating the mean latent coordinates of particles in the latent space which originated from each batch, the average path of the MD trajectory through the latent space embedding was traced. The gen-To evaluate the decoding of the latent space into density maps we compared the 50 generated maps to 50 atomic models at equidistant time intervals in the MD trajectory by real-space map-to-model correlation and plotted those in a 2D heatmap shown in Fig. 5C. The heatmap features a strong diagonal, indicating that the sampled volumes are most similar to the conformation of the molecule in the particles around the selected latent coordinate. We highlight a few sampled density maps together with the atomic model which has the highest correlation to that map in Fig. 5D. A more detailed map-to-model fit is shown erated path progresses continuously through the latent space embedding. We then used the trained decoder to evaluate these selected latent coordinates and produce their corresponding 3D density maps.

**Figure 5.**
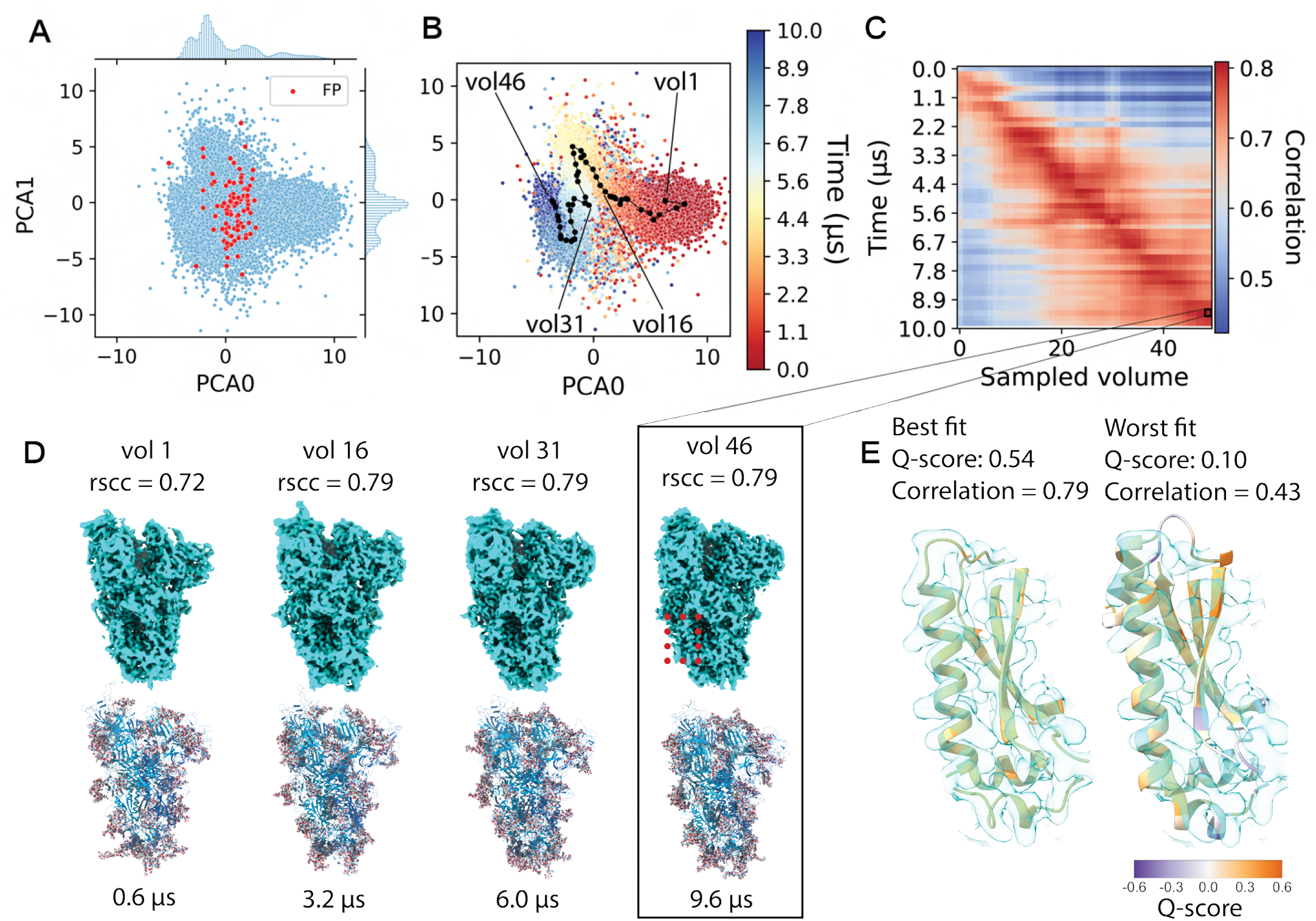
Analysis of heterogeneous reconstruction of SARS-CoV-2 spike trimer. **A**: Plot of the latent space after applying PCA. Red points indicate the false positive particles in the latent space. **B**: Each particle embedding is coloured according to the frame of the MD trajectory the particle originated from. Black dots show a trajectory traced through the latent space. **C**: Real-space correlation between 50 volumes generated by sampling the latent space (x-axis) and frames in the MD trajectory (y-axis). **D**: Example volumes sampled from the latent space with the frame of the trajectory for which it showed the highest correlation. **E**: Zoom of the density map of volume 46 with the best and worst atomic models out of the 50 sampled states from the MD trajectory displayed. Atomic models are coloured by backbone Q-scores.

in Fig. 5E where volume 46 is shown with a close-up of the best and worst fitting atomic models. We further validated the map-to-model fit by calculating the average Q-scores (Pintilie *et al*., 2020) for both models and found a Q-score of 0.54 for the best fitting model compared to a score of 0.10 for the worst fitting model. Atomic structures are coloured according to backbone per-residue Q-score. Entire atomic models coloured by backbone per-residue Q-score are shown in supplementary Fig. S4B and C.

In addition, we trained CryoDRGN on a similar dataset where radiation damage was enabled, as described in section 2.3. The latent space is plotted in Fig. S5. We again found that CryoDRGN was able to organise the particles according to their timepoint in the MD trajectory, despite the decreased SNR of the high-frequency information and the comparatively small number of particles (15846) used to train CryoDRGN in this case.

Next, we repeated the same steps for a second dataset based on the SARS-CoV-2 RTC. The MD simulation that was used as a source of heterogeneity for this synthetic dataset is much longer (500 µs compared to 10 µs for the spike protein) and sampled with a time-interval of 4.8 ns (Shaw, 2020). From this trajectory 10000 frames were sampled and 167 micrographs were simulated with a total of 50100 particles. Fig. 6A shows the consensus reconstruction obtained with 41146 particles, which reached 3.11 Å global resolution. Despite the much longer simulation, CryoDRGN again was able to organise the latent space coherently with timepoints in the MD trajectory from which the particles were sampled, as exemplified in Fig. 6B and C. We used the same approach as to the open-state SARS-CoV-2 dataset to compute a real-space correlation matrix, which is shown in Fig. 6D. Good agreement was found between the timepoints in the MD trajectory and the generated volumes, although towards the later timepoints in the trajectory we see broader correlations with the volumes generated. This may indicate that conformations become less distinguishable. This conclusion is supported by the averaged path through the latent space in Fig. 6C, and the lack of separation in the latent space between later timepoints in the MD simulation is easily observed in the high density region in Fig. S4D. Finally, we show two examples of a small section of the atomic model in a flexible region of the complex in Fig. 6E; the best and worst fit according to real-space map-to-model correlation. The entire atomic model coloured by backbone Q-scores is shown in Fig. S4E and F.

**Figure 6.**
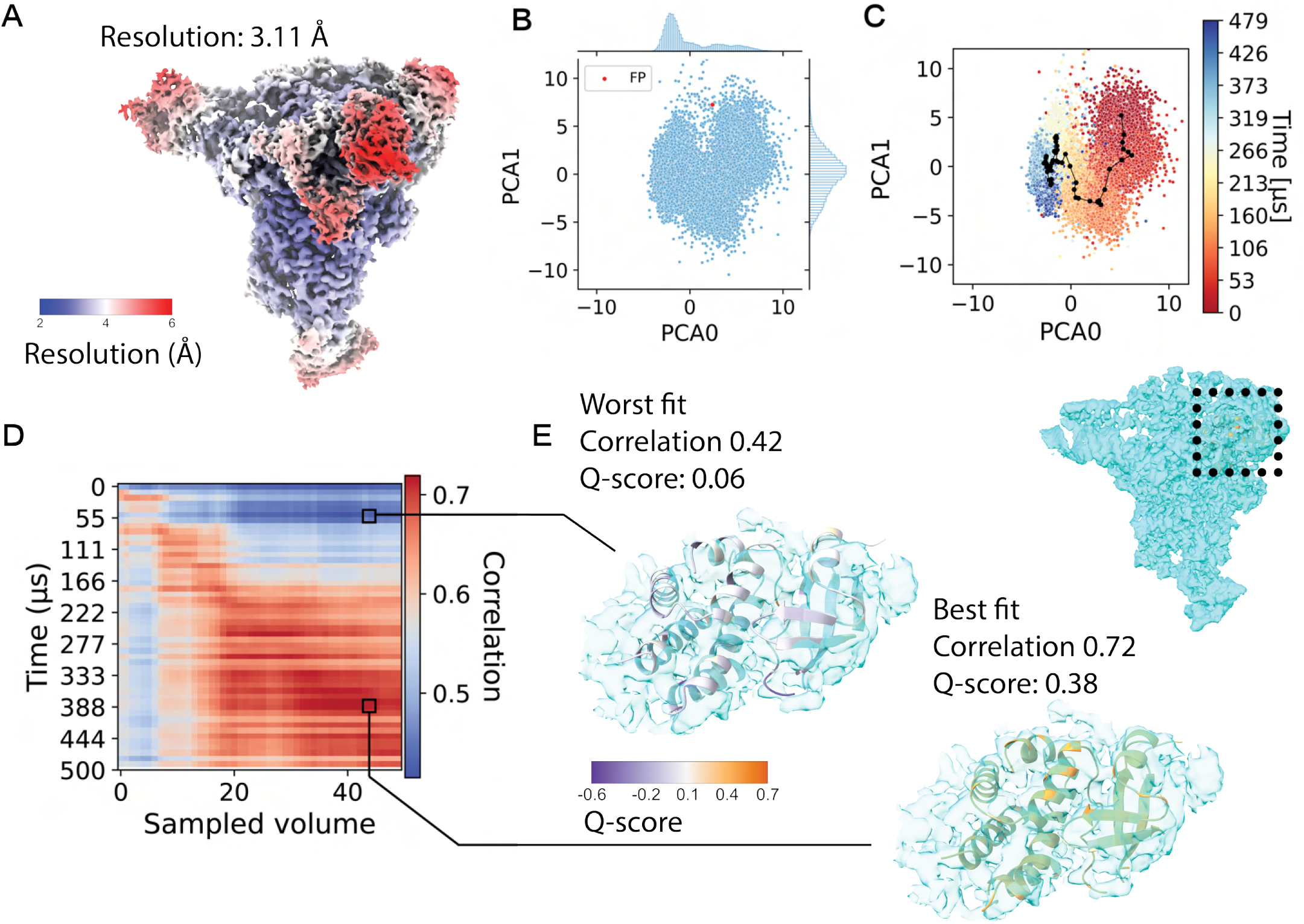
Analysis of heterogeneous reconstruction of the SARS-CoV-2 RTC. **A**: Consensus reconstruction of the RTC, coloured by local resolution. **B**: Plot of the latent space after applying PCA. **C**: Same latent space, coloured according to the frame of the MD trajectory the particle originated from. **D**: Real-space correlation between 50 volumes generated by sampling the latent space (x-axis) and frames in the MD trajectory (y-axis). **E**: Best and worst atomic models shown in the density map of volume 43.

As discussed in section 2.2 we also performed a steered MD simulation of the SARS-CoV-2 spike protein. We reduced the complex to a single monomer and forced a conformational change between an open and closed state of the RBD using harmonic restraints. We then trained CryoSPARC’s 3DFlex model using 1-dimensional and 2-dimensional latent spaces. The latent spaces of both trained models are shown in Fig. S6A-D (for the 1-dimensional latent space we show the distribution of latent coordinates). In the case of a 1-dimensional latent space we find that there is some correlation between the estimated latent encoding and the timepoint in the MD trajectory, whereas in the 2-dimensional latent space we find that the latent space is not organised in a continuous manner. Unlike our previous examples, there is no path through the cluster that can reconstruct the MD trajectory in chronological order. Correlation between the timepoints in the MD trajectory and the sampled volumes, shown in Fig. S6E and F show a strong relation between conformations in the MD trajectory and the sampled volumes in the case of a 1-dimensional latent space, but not for a 2-dimensional latent space.

We tested 3DFlex with another synthetic dataset based on a steered MD simulation of the complement system protein C3, based on a morph trajectory interpolating between PDB 2a73 and PDB 2i07. C3 undergoes a major conformational change when transitioning from its inactive state C3 to the active C3b that exposes a reactive thioester for opsonisation of target surfaces (Janssen *et al*., 2006). A visualisation of the ensemble of states generated during the steered MD simulation is shown in Fig. S7A and a consensus reconstruction from 96169 particles with 2.55 Å global resolution in Fig. S7B. We then trained a 3DFlex model with a two-dimensional latent space on this data and obtained a latent space where different sections of the MD trajectory were clustered together. Due to the magnitude of the conformational change we ran the steered MD simulation in four iterations with the target state first set to a conformation a third of the way between the initial and final state, then being changed to a state two-thirds of the way and then being changed to the final state for the last two iterations. As a result, we expected that the latent space might be split into three discrete clusters that resemble the target states. We observe three regions in the latent embedding corresponding to the first 45 ps, the next 45-90ps and the last 110ps timepoints. However, there is not a continuous path through the latent space following the chronological order of the trajectory and these regions are not well separated.

## 3. Discussion

Many biomolecules have evolved to perform specific tasks through a concerted sequence of conformational motions. The current trend in SPA is to address conformational heterogeneity with machine learning algorithms in order to identify and sort a continuum of structural states into a continuous representation of heterogeneity which may be correlated to functional motion. Here, we have demonstrated a new toolkit for the simulation of single-particle cryo-EM micrographs that contain conformational heterogeneity, investigated the effect of this heterogeneity on consensus reconstructions and explored the ability of two established machine learning-based HRAs to quantitatively recover the ground-truth conformational heterogeneity from a collection of single-particle snapshots.

We found that training both CryoDRGN and 3DFlex resulted in models for which particles with conformations that were temporally close in the MD trajectory were also clustered closely together in the latent space. Using CryoDRGN to process the SARS-CoV-2 spike glycoprotein and RTC synthetic datasets we found that it was possible to generate a path through the latent space such that the reconstructed volumes correlated strongly with evenly spaced timepoints of the MD trajectory.

The ultimate goal would be to decode the chronological order of states underlying functionally relevant conformational trajectories. However, at present we focus on structures that are conformationally similar during traversal through latent space despite the non-linear embedding. One area of future study could be devoted to developing and evaluating methods to interpret these learned latent spaces. One such example is CryoDRGN conformational landscape analysis (Zhong, 2022) which, through analysis in volume-space, both clusters a small number of discrete conformational states and allows inference of continuous reaction coordinates. With the presence of ground truth information, investigating the optimisation of dimensionality reduction and clustering methods applied to both MD trajectories and latent spaces may provide insights on the interpretation of both.

Our analysis also highlights that more work is needed to understand the requirements on data quality for HRAs to recover physically relevant conformational trajectories. In this paper we limited our investigation to a coarse analysis of the effects of radiation damage and fluence on conventional reconstructions, but it is possible to conduct systematic studies into the behaviour of HRAs in the limit of poor data quality.

Other aspects of cryo-EM image processing workflows can also be investigated using this approach. We have demonstrated this for particle picking by using the ground-truth particle positions to compute precision and recall for a set of picked particles. As this aspect of the pipeline often involves parameter optimisation as well, it may be beneficial to investigate the behaviour of particle pickers, 2D/3D classification algorithms and refinement algorithms in the limit of strong conformational or compositional heterogeneity, very poor SNR, unevenly sampled particle orientations or poor annotations of training data. It is also possible to simulate experimental pathologies such as preferred orientation and to investigate the effect this has on conventional 3D reconstruction workflows or HRAs. Conversely, Roodmus also makes it possible to study the effects which future developments in instrumentation and algorithms may have on the ability to reconstruct heterogeneity.

Roodmus sources conformational heterogeneity from MD simulations. This presents a major advantage over previous, related methods where the conformational heterogeneity was generated by interpolating between two or more discrete structural states. Atomistic ensembles from MD simulations allow evaluation of HRAs in more physically realistic settings, but come with the disadvantage that large-scale domain motion is computationally expensive to simulate at the atomistic level. Coarse-grained MD or enhanced sampling strategies may be a solution that could allow for more in-depth exploration of the behaviour of both conventional reconstruction workflows and HRAs in the case of large-scale motion, or in cases where intermediate states are less densely populated.

Image simulation with accurate modelling of experimental image contrast and optical aberrations is essential to evaluate HRA performance on realistic cryo-EM datasets with independent ground-truth information. The multislice forward model of the Parakeet software offers improved cryo-EM image simulation methods compared to alternative ways of generating synthetic data. An electron wave is propagated through a virtual sample, as opposed to a simple linear projection through the volume. This more accurately simulates the projection of the specimen’s electrostatic potential. Parakeet included models for detector response, contrast modulation by the CTF and a sophisticated model for structural noise contributions from an amorphous ice layer. We found no qualitative effect of Parakeet’s radiation damage model on the training of CryoDRGN and we still obtained consensus reconstructions with unusually low B-factors compared to experimental data. This suggests refinement of the model i s required to more accurately reflect the structural damage inflicted o n t he s pecimen d uring electron exposure, which represents one of the main limiting factors of cryo-EM imaging (Hayward & Glaeser, 1979). An additional missing element in Parakeet simulations is that of beam-induced motion. Since such motion can only be imperfectly corrected for, including these effects would likely yield overall B-factors and concomitant attenuation of high-frequency signal more similar to those observed in experimental data. Synthetic datasets that feature these properties may be more realistic starting points for investigating the data quality requirements of HRAs.

As current HRA methods evolve and improve, periodic community evaluation of these methods against ground truth data using the Roodmus toolkit would help quantify the performance of these methods and encourage wider usage in the field. Further comparisons could also be made to orthogonal experimental data for the same (or similar) biological systems. Comparison of simulated and experimental datasets may help understand the degree to which molecular dynamics trajectories underestimate experimental disorder given their simulation time.

Roodmus (Greer *et al*., 2024) is available as open source software (github.com/ccpem/roodmus) and from PyPI (pypi.org/project/roodmus). The Roodmus toolkit provides the necessary modular utilities to generate synthetic SPA datasets and explore the performance of processing algorithms on them. Whilst it is important to be cogniscient that Roodmus can only be used to explore heterogeneity within the bounds of the MD simulation data provided to it, and that there are avenues for improvement in the simulation protocol, we believe that Roodmus facilitates the undertaking of a range of studies for elucidation of conformational heterogeneity investigation techniques. We have reported findings from studies utilising both discrete and continuous approaches to exploring heterogeneity on a number of synthetic datasets but there is ample opportunity for further work to better understand conventional reconstruction pipelines, the data quality requirements of HRAs and the traversal and clustering of latent encodings of heterogeneity. There is also scope to investigate the relation between synthetic and experimental SPA datasets, methods for improvement of synthetic micrograph generation and the extension of Roodmus to tomography.

## 4. Methods

### 4.1. MD trajectories

Publicly available MD trajectories of the SARS-CoV-2 RTC and the spike glycoprotein in closed and partially open states were performed by (Shaw, 2020) and downloaded from https://www.deshawresearch.com Specific trajectories were DESRES-ANTON-13795965 for the RTC, DESRES-ANTON-[11021566,11021571] for spike protein in the closed and partially open states respectively.

Additional steered MD trajectories were produced using the openMM (Eastman *et al*., 2017) Python library. PDB models 6xm4 and 6xm5 (Zhou *et al*., 2020) were used as starting and target conformation respectively. Chain B was isolated from both models and prepared for simulation using the pdbfixer library provided by openMM. Harmonic restraints were added between each pair of C-alpha atoms in the starting and target models. The system was calibrated at 0 K for 50 ps and then annealed to 300 K for another 50 ps. The force constant for all restraints was annealed between 100 and 400 ps from 0 to 10 kJ nm^−1^ mol^−1^. The integration time step was 2 fs, and the time interval between saving frames was 10 fs. After completion of the simulation, the trajectory was downsampled to 10000 contiguous frames between 100 and 200 ps in which the molecule underwent the conformational change.

A steered MD simulation was performed on human complement component C3 (PDB ID 2a73; (Janssen *et al*., 2006)) as it undergoes a conformational transition into a state termed C3b (PDB ID 2i07; (Janssen *et al*., 2006)). A morphing trajectory interpolating between both states was kindly provided by F. Forneris (University of Pavia). Starting from frame 1 of the morph and taking frame 31 as a target, similar harmonic restraints as described above were added between each pair of C-alpha atoms in the structures. An integration timestep of 2 fs was again used with a time interval between saved frames of 100 fs. The system was initialised at 300 K with a force constant of 5 kJ nm^−1^ mol^−1^, which was annealed to 15 kJ nm^−1^ mol^−1^ over 200 ps. After 200 ps, the target structure was replaced with frame 61 of the morph and the force constant was annealed again from 5 to 15 kJ nm^−1^ mol^−1^ over 200 ps. Another 400 ps were simulated with frame 91 as the target state, first 200 ps with a force constant increasing from 10 to 60 kJ nm^−1^ mol^−1^, then another 200 ps with a force constant increasing from 60 to 120 kJ nm^−1^ mol^−1^.

### 4.2. Synthetic micrograph and movie simulation

All image simulation was performed using Parakeet software of git commit 17a0c864f6cfd84b5fd56b60fa446f7b021d338c available from github.com/rosalindfranklininstitute/parakeet. The Roodmus *conformations sampling* utility was used to sample a number of conformations from each MD trajectory, as indicated in table 1. Roodmus *run parakeet* utility was used to configure and run Parakeet to produce micrographs containing 300 particles (SARS-CoV-2 spike trimer and RTC) or 250 particles (steered MD datasets).

**Table 1.** Specification of simulated datasets and refinement statistics.

|  | SARS-CoV2<br>spike trimer (closed) | SARS-CoV2<br>spike trimer (open) | SARS-CoV2<br>spike trimer (mixed) | SARS-CoV2<br>RTC | SARS-CoV2<br>spike monomer | C3-C3b |
| --- | --- | --- | --- | --- | --- | --- |
| Trajectories | DESRES-ANTON-11021566 | DESRES-ANTON-11021571 | DESRES-ANTON-[11021566, 11021571] | DESRES-ANTON-13795965 | 6xm4-6xm5 steered | 2a73-2i07 steered |
| # of conformations | 8,334 | 8,334 | 8,334 (x2) | 10,000 | 10,000 | 2,000 |
| # of particles | 270,000 | 270,000 | 120,000 | 50,100 | 200,000 | 200,000 |
| Acceleration voltage (kV) | 300 | 300 | 300 | 300 | 300 | 300 |
| Fluence ($e^-/\text{\AA}^2$) | 45 | 45 | 45 | 45 | 45 | 45 |
| Flux density ($e^-/\text{pixel/s}$) | 5 | 5 | 5 | 5 | 5 | 5 |
| Pixel size ($\text{\AA}$) | 1.0 | 1.0 | 1.0 | 1.0 | 1.0 | 1.0 |
| Defocus range ($\mu\text{m}$) | 0.5 - 3.5 | 0.5 - 3.5 | 0.5 - 2.5 | 0.5 - 3.5 | 0.5 - 2.5 | 0.5 - 2.5 |
| Refinement software | RELION 4.0 | RELION 4.0 | RELION 4.0 | RELION 4.0 | CryoSPARC 4.2.1 | CryoSPARC 4.2.1 |

The RTC dataset was generated as movies with 3 frames with no particle motion between frames and no radiation damage simulation. The movies with beam-damage enabled, based on MD trajectory DESRES-ANTON-11021571 introduced in section 2.3, were generated with 30 frames and a total fluence of 45 *e*^−^/Å^2^, allowing a sensible use of the Parakeet radiation damage model (Parkhurst *et al*., 2021).

All datasets except those noted in section 2.3 were simulated at 45 electrons per square angstrom, with a pixel size of 1.0 Å and an ice thickness of 50 nm which is typical for many single-particle datasets (Noble *et al*., 2018). For lower fluence datasets, 100 micrographs were simulated with 30000 total particles per condition. Fluence parameters were varied between 45 and 5 *e*^−^/Å^2^.

Simulation of datasets was benchmarked utilising an Intel Xeon Gold 5218 processor with 75GB of RAM and Nvidia Tesla V100 (32GB) GPU. Generating a single non-fractionated micrograph following the simulation parameters used for the spike trimer (open) dataset took 5 minutes. Generating a single 30-frame fractionated movie with radiation damage took 22 minutes. We note that simulation time depends on a number of simulation parameters, including the number of particles, fractionation, sample size and which physical effects are simulated.

### 4.3. cryo-EM processing in RELION

The datasets created from the DESRES-ANTON-13795965 and DESRES-ANTON-[11021566,11021571] MD simulations were reconstructed using RELION 4.0. Altogether, these constitute: the single-micrograph datasets for DESRES-ANTON-[11021566,11021571], the DESRES-ANTON-11021571 dataset utilising a single conformation, the 30-frame fractionated movie dataset for DESRES-ANTON-11021571, the 3-frame movie dataset for DESRES-ANTON-13795965 and the DESRES-ANTON-[11021566,11021571] mixed dataset. The workflows for consensus reconstructions of these are illustrated in Figs. S8, S9, S10 and S11. along with visualisations of several stages of the processing. Depicted micrographs were normalised with the ccpem-pipeliner gitlab.com/ccpem/ccpempipeliner.

Before CTF estimation using CTFFIND4 (Rohou & Grigorieff, 2015), the movie datasets were motion-corrected -an unnecessary step for the single-micrograph datasets. Topaz (Bepler *et al*., 2019) was trained on a manually picked subset of particles and used for autopicking. 100 2D classes were generated and those with a *relion class ranker* score greater than 0.25 were kept. After initial model building, 4 3D classes were deduced and poor classes were removed after manual inspection. Those remaining were 3D refined to produce a single consensus map suitable as the input for a PostProcess job (to determine the global resolution after applying a mask) and for local resolution determination via a LocalRes job. The number of particles kept at each stage of reconstruction is reported in Figs. S8 to S11.

### 4.4. cryo-EM processing in CryoSPARC

The SARS-CoV-2 spike monomer steered MD dataset and the C3-C3b steered monomer datasets were reconstructed using CryoSPARC 4.2.1. For the monomeric SARS-CoV-2 spike dataset 800 micrographs were imported. CTF estimation was performed using CTFFIND4, followed by automatic particle picking with the blob picker algorithm, with minimum and maximum diameters of 50 and 200 Å, resulting in 1968039 picked particle locations. Picked particles were filtered based on the local power, extracted and 2D classified into 50 classes. A selection of 19 classes (93574 particles) was then used for Topaz training, resulting in 368967 particle picks. These picked particles were again filtered, extracted and 2D classified into 50 classes. All classes (213486 particles) were selected for Ab-Initio model building with 4 classes. The first class was selected as a reference for homogeneous refinement with all 213486 particles. This consensus reconstruction was then used to train 3DFlex as described in section 4.6. The workflows for consensus reconstruction of these datasets are illustrated in Fig. S12 and Fig. S13 along with visualisations of several stages of the processing.

For the C3-C3b steered MD dataset 800 micrographs were imported. CTF estimation was performed using the Patch CTF estimation job, followed by manual picking of 267 particles in 4 micrographs. These picked particles were used to train Topaz, resulting in 183092 picked particle locations. The picked particle locations were filtered, extracted and 2D classified into 50 classes. All particles were kept, resulting in 96169 particles used for ab-initio model building with 4 classes. The second class was used as a reference for homogeneous refinement with all particles. This consensus reconstruction was then used to train 3DFlex as described in section 4.6.

### 4.5. Training CryoDRGN

Pose and CTF information obtained from the consensus reconstruction in RELION were extracted using cryoDRGN version 3.0.0b0 according to the CryoDRGN tutorial (ezlab.gitbook.io/cryodrgn). The model was then trained for 25 epochs with default residual MLP architecture for both the encoder and decoder using particle images of 320×320 pixels (1 Å pixel size). The dimensionality of the latent space was 8 in all cases. The number of particles used for training were 41146 for the SARS-CoV-2 RTC dataset (DESRES-ANTON-13795965), 236079 for the SARS-CoV-2 spike glycoprotein dataset (DESRES-ANTON-11021571) and 15846 for the dose-fractionated dataset.

### 4.6. Training 3DFlex

Training the 3DFlex model was done according to the tutorial provided by CryoSPARC. Particles taken from the homogeneous refinement were used for a 3D Flex Data Prep job with a training box size of 128 pixels. A mesh was created with a base number of tetrahedral cells of 30. The model was trained with a latent dimensionality of 2, 64 hidden units in the flow generator network for 24 epochs. Custom trajectories through the latent space were created with the cryosparc-tools Python library.

### 4.7. Dataset availability

Synthetic micrographs and atomic structure models used in data generation, as well as reconstructed density maps and image processing intermediate results are available from 4TU.ResearchData repository.

Data can be downloaded for DESRES-Trajectory sarscov2-11021571-all-glueCA single conformation dataset:

https://data.4tu.nl/private_datasets/ugJkrhdbTnXLXc4CD1PlHtob0JcD071lQFKWnidA0UY.

Data can be downloaded for DESRES-Trajectory sarscov2-11021571-all-glueCA:

https://data.4tu.nl/private_datasets/mNkVIFZj5FaJh95sr6W7MSA03ppPIOA63W2gvAlgehA.

Data can be downloaded for DESRES-Trajectory sarscov2-11021571-all-glueCA single fractionated:

https://data.4tu.nl/private_datasets/I1q-quITgg-o2m68JawJ04MiKOMnwyjxeFYOD6JAFaM.

Data can be downloaded for DESRES-Trajectory sarscov2-11021566-11021571-mixed:

https://data.4tu.nl/private_datasets/vT_

Jsu3GSPX_G4Vu_08oymMv6vYt514lSPT9wwf8bLE.

Data can be downloaded for DESRES-Trajectory sarscov2-13795965-no-water-movies:

https://data.4tu.nl/private_datasets/jCOtSqXp52aCoI-ES10AGsyoVFtqXvclEmRY3jSCO8A.

Data can be downloaded for 6xm5 steered:

https://data.4tu.nl/private_datasets/SUpnL6U_PbJiPkHsX5e6GrPHOPtuqH48vkfE916jZ8c.

Data can be downloaded for c3c3b:

https://data.4tu.nl/private_datasets/9m3Q20VSvfsxjcYKUDWMwzmLE-yjQQQ27cYwAoqnopI.

Image processing results and intermediates from RELION and CryoSPARC can be downloaded for all datasets from: https://data.4tu.nl/private_datasets/v1UdxmIX5B-QeNhjKClFltNk7xP07KuCVFdLPFUTU9k.

## Acknowledgements

MJ and AJ thank the TU Delft AI initiative and the Kavli Institute for Nanoscience Delft for financial support, and acknowledge the use of computational resources of the DelftBlue supercomputer provided by the Delft High Performance Computing Centre. JG and TB thank MRC Partnership Grant MR/V000403/1; Wave 1 of The UKRI Strategic Priorities Fund under the EPSRC Grant EP/W006022/1, particularly the “AI for Science” theme within that grant & The Alan Turing Institute; and support from the Ada Lovelace Centre. We thank Federico Forneris for providing the initial morph trajectory used for the steered MD simulation of C3-C3b.

## Supplementary Figures

**Figure S1.**
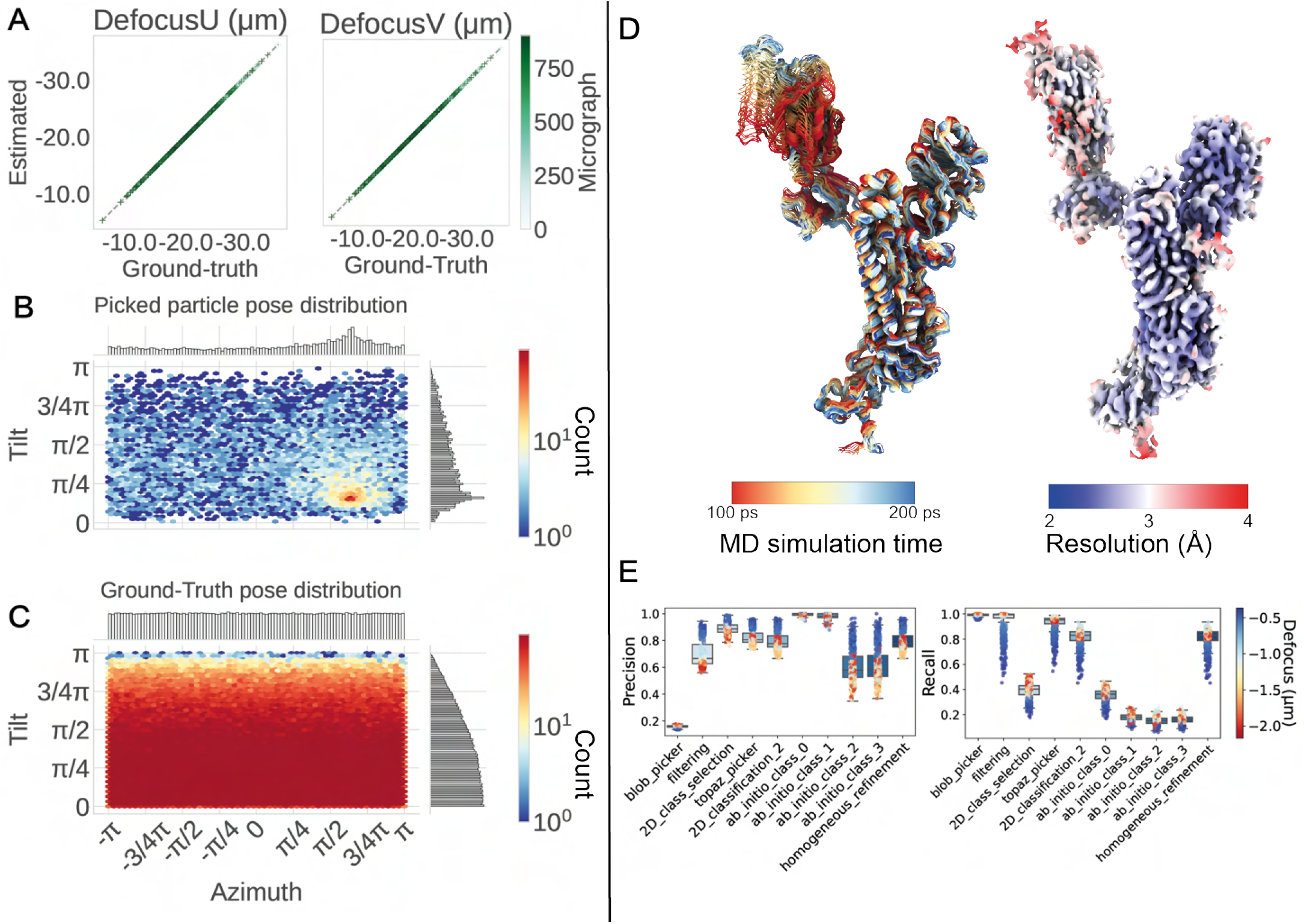
Further processing statistics for spike trimer protein and reconstruction of spike monomer. **A**: Correlation between estimated and true defocus values for each micrograph in heterogeneous spike trimer (open) dataset. **B**: Distribution of orientations estimated during 3D refinement of heterogeneous spike trimer (open) 8000 particle subset. **C**: True orientations for each particle in total spike trimer (open) dataset. **D**: Monomer SARS-CoV-2 spike protein, model and reconstructed consensus density map, colour indicates local resolution. **E**: Particle picking precision and recall for spike monomer. X-axis indicates job types used during processing in CryoSPARC.

**Figure S2.**
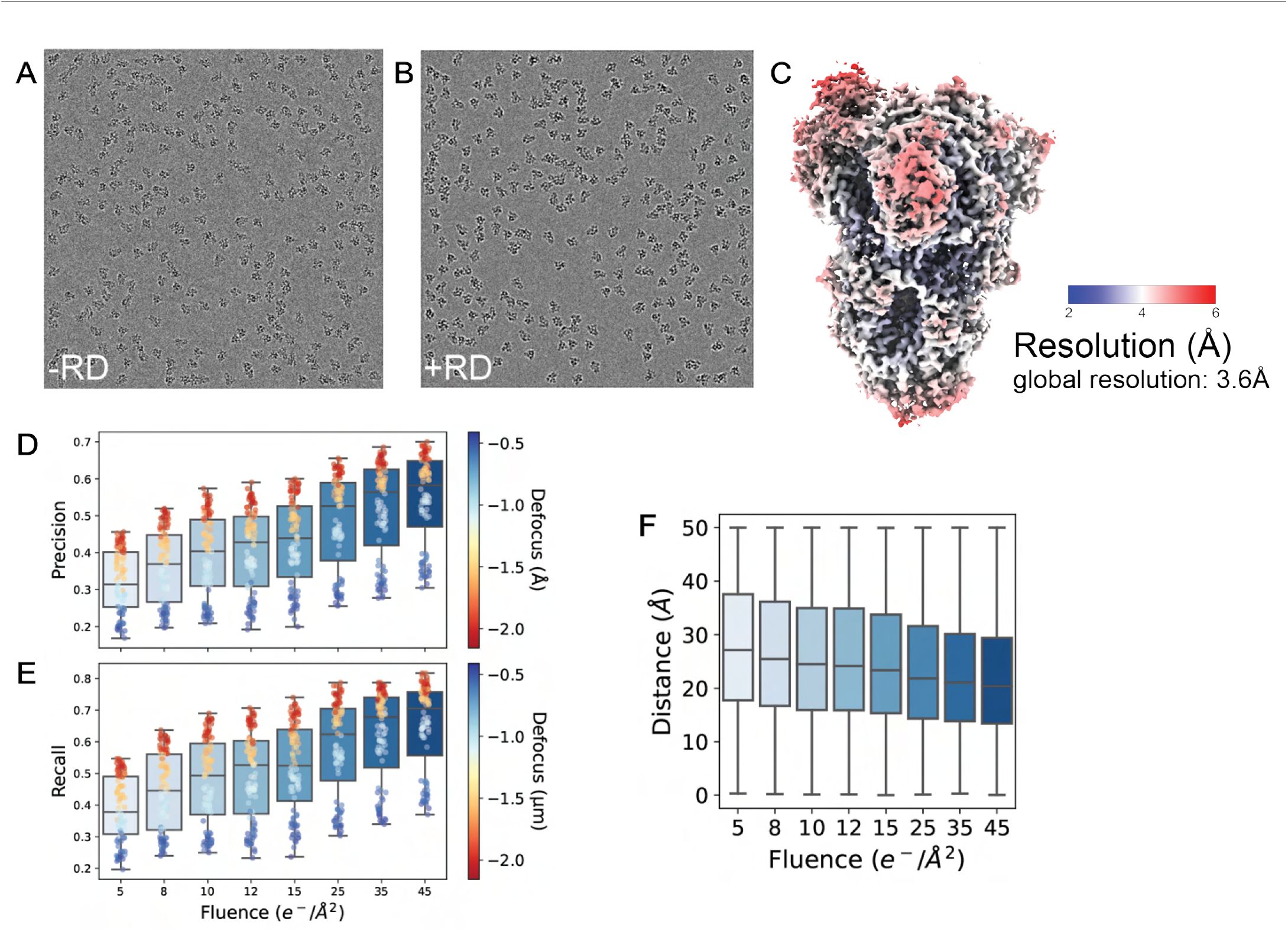
Effect of defocus and fluence on synthetic micrographs. **A**,**B**: Example micrograph shown after simulation without (**A**) and with (**B**) radiation damage. Normalisation for visualisation done with the ccp-em pipeliner. **C**: Reconstructed consensus density map of the dataset with radiation damage, using 15846 particles. **D**,**E**: Picked particle precision and recall for datasets simulated with increasing total fluence. **F**: Boxplot showing the distribution of distances between each picked particle and the closest ground-truth particle within 50 Å . The median decreases as fluence increases, indicating that the picked position becomes more accurate w.r.t. the ground-truth position as the fluence increases.

**Figure S3.**
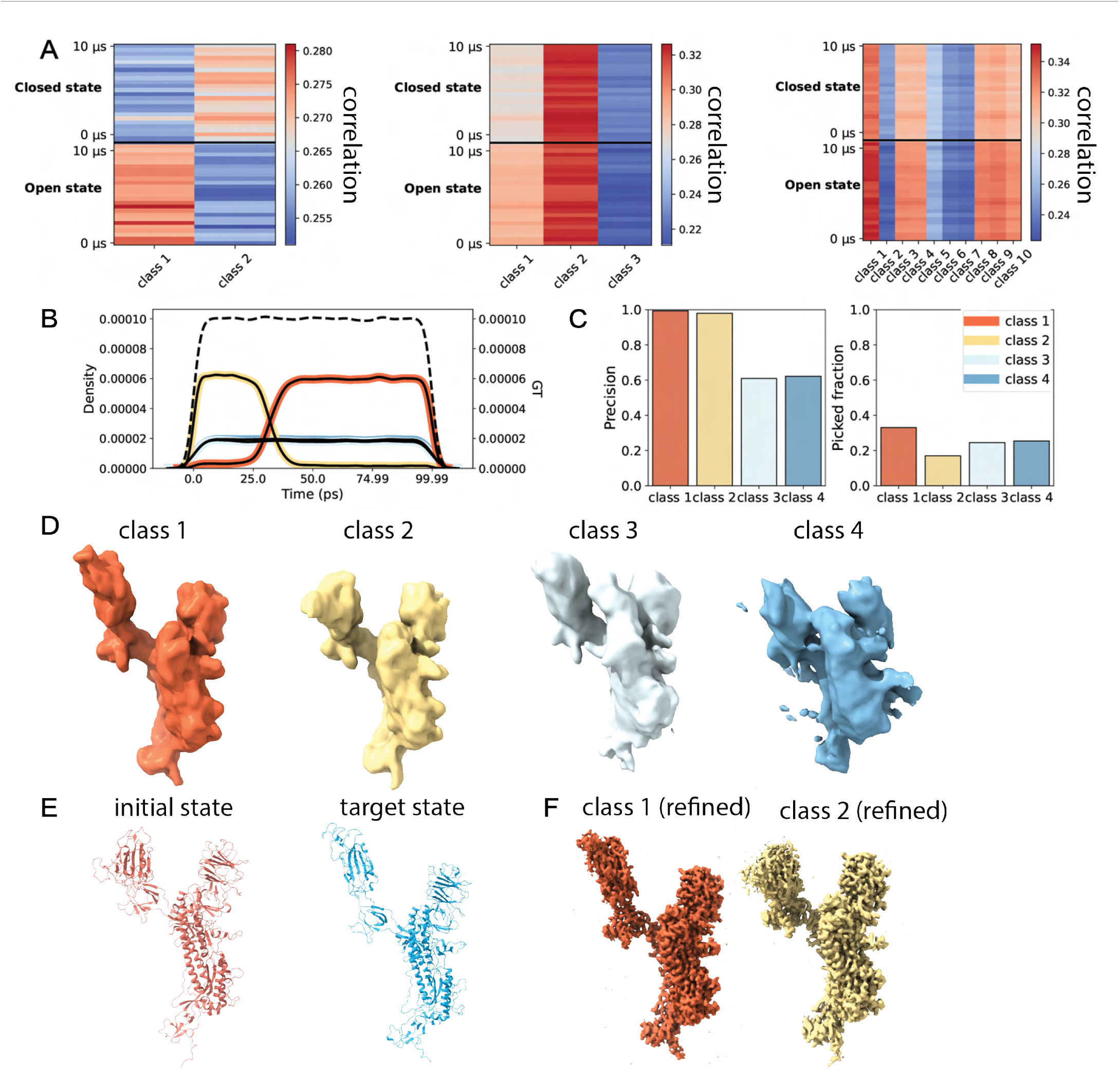
3D classification of steered MD dataset of SARS-CoV-2 spike monomer. **A**: Unnormalised correlation heatmaps for the 2, 3 and 10-class classification of the SARS-CoV-2 spike glycoprotein mixed dataset. For each class the real-space cross-correlation was computed with 25 evenly spaced models from the closed state and the open state MD trajectories. **B**: Distribution of particles in each class over the frames of the MD trajectory. **C**: Precision per 3D class (left) and fraction of particles in each 3D class (right). **D**: Initial models of each 3D class, only class 1 and 2 were selected for refinement. **E**: Atomic models of the first and last conformation of the MD trajectory respectively. **F**: Refined density maps for class 1 and 2. Visual inspection shows that class 1 resembles the target conformation, while class 2 resembles the initial conformation.

**Figure S4.**
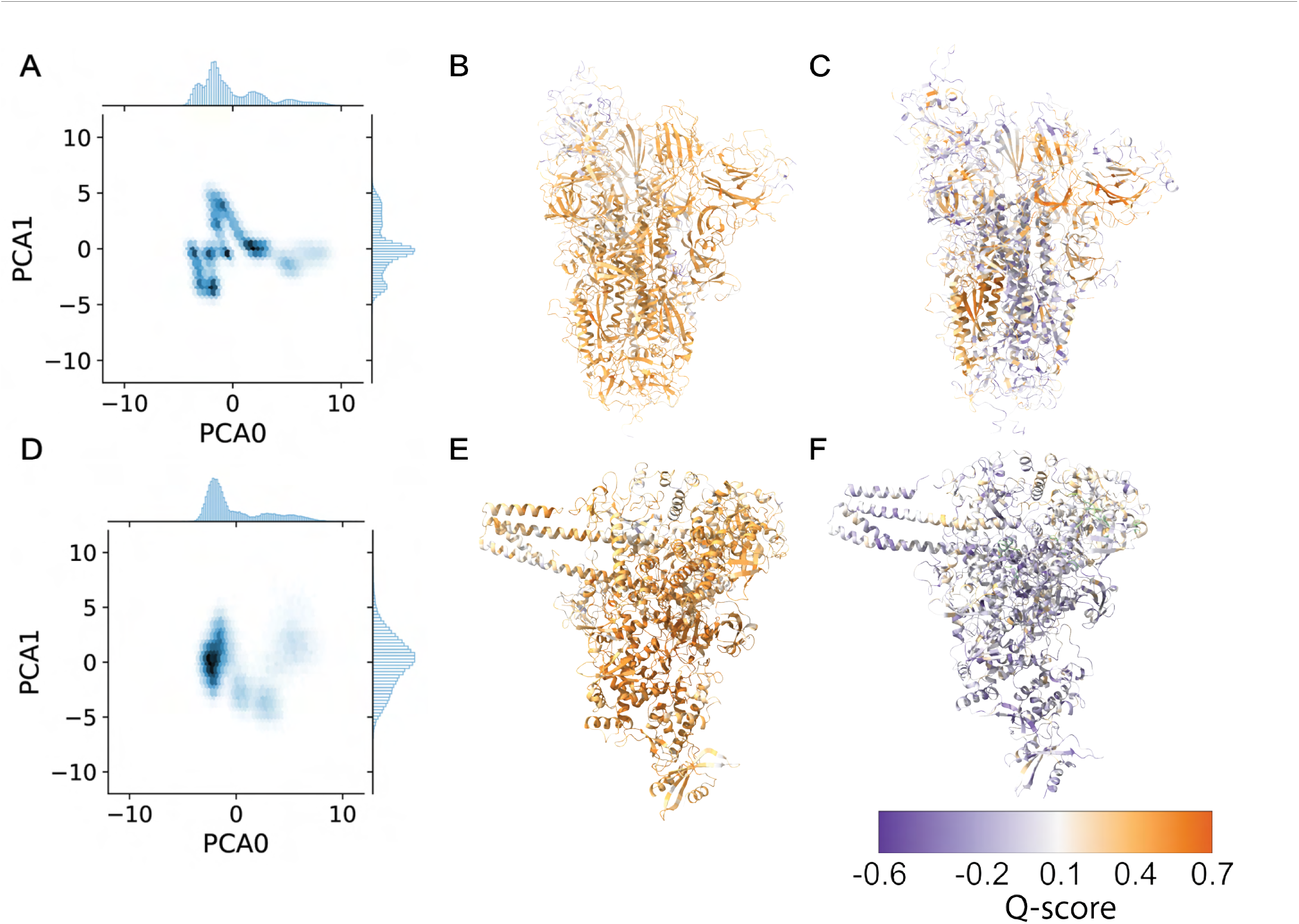
Further analysis of heterogeneous reconstruction of SARS-CoV-2 spike trimer and RTC.. **A**: Latent space of CryoDRGN training, visualised as a hexbin plot. This better shows the density in each region of the latent space. **B**,**C**: Best and worst atomic model fit to sampled volume 46 of spike protein CryoDRGN training. **D**: Hexbin visualisation of the latent space of the CryoDRGN training done for the RTC dataset. **E**,**F**: Best and worst atomic model fit to sampled volume 43 for RTC complex CryoDRGN training.

**Figure S5.**
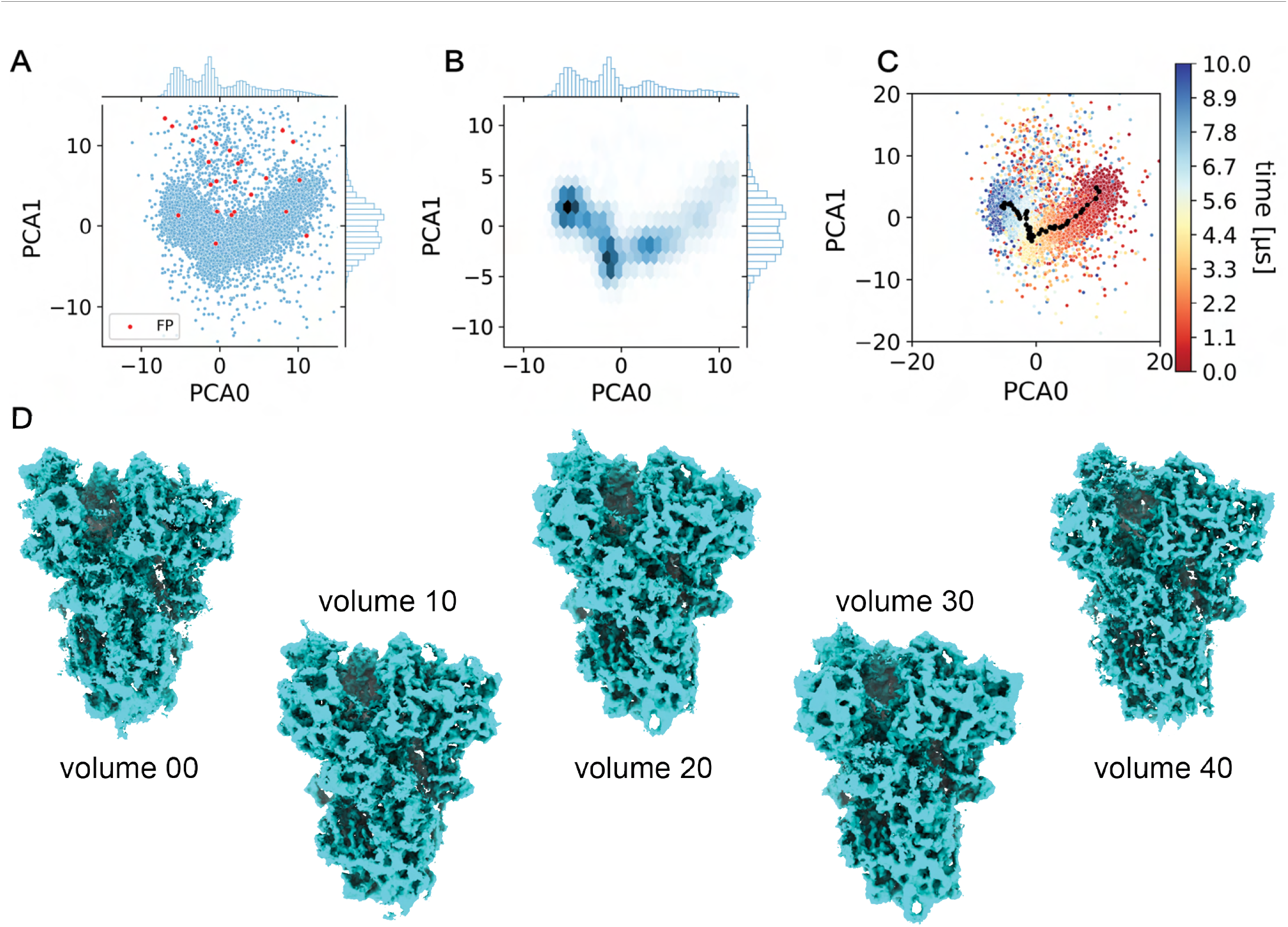
CryoDRGN training on the dose-fractionated dataset. **A**: Scatterplot of the latent space, with false positive (FP) particles marked in red. **B**: Hexbin plot of the latent space, showing it consists of a continuous region of density. **C**: Latent space coloured by timepoint in the MD trajectory. **D**: Example volumes generated from the averaged latent space coordinates in **C**.

**Figure S6.**
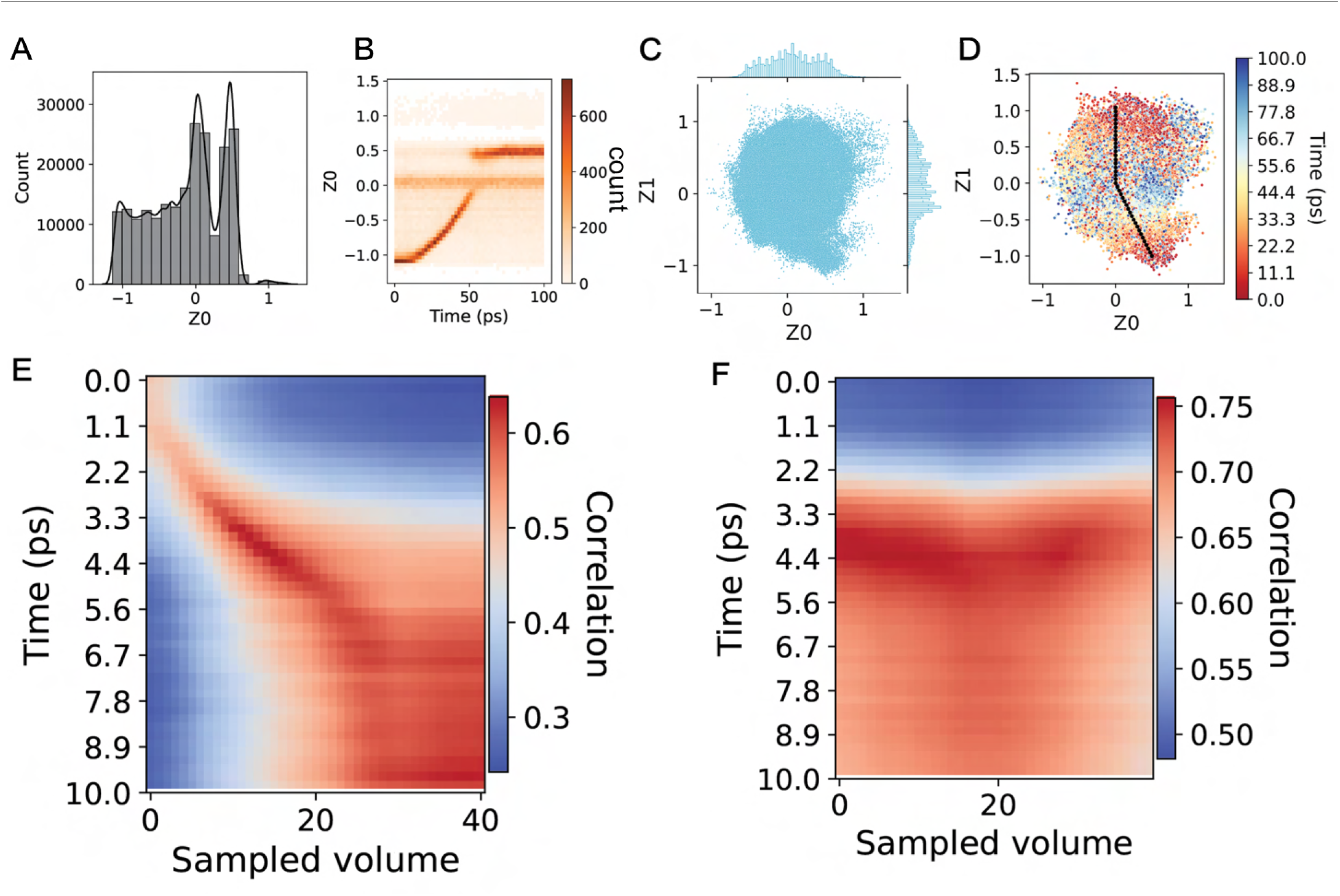
3DFlex training for the SARS-CoV-2 spike monomer with zdim = 1 and zdim = 2. **A**: Distribution of latent space coordinates (zdim = 1). **B**: 2D histogram of the latent space coordinates and the timepoint in the MD trajectory of each particle. This plot shows a strong relation between the latent space coordinates and the trajectory. **C**: Latent space from training with zdim = 2. **D**: Latent space where each particle is coloured by its corresponding timepoint in the MD trajectory. Traversal through the latent space is plotted in black. **E**: Real-space correlation heatmap between the MD trajectory and volumes sampled from the 1-dimensional latent space. **F**: Real-space correlation heatmap for the 2-dimensional latent space.

**Figure S7.**
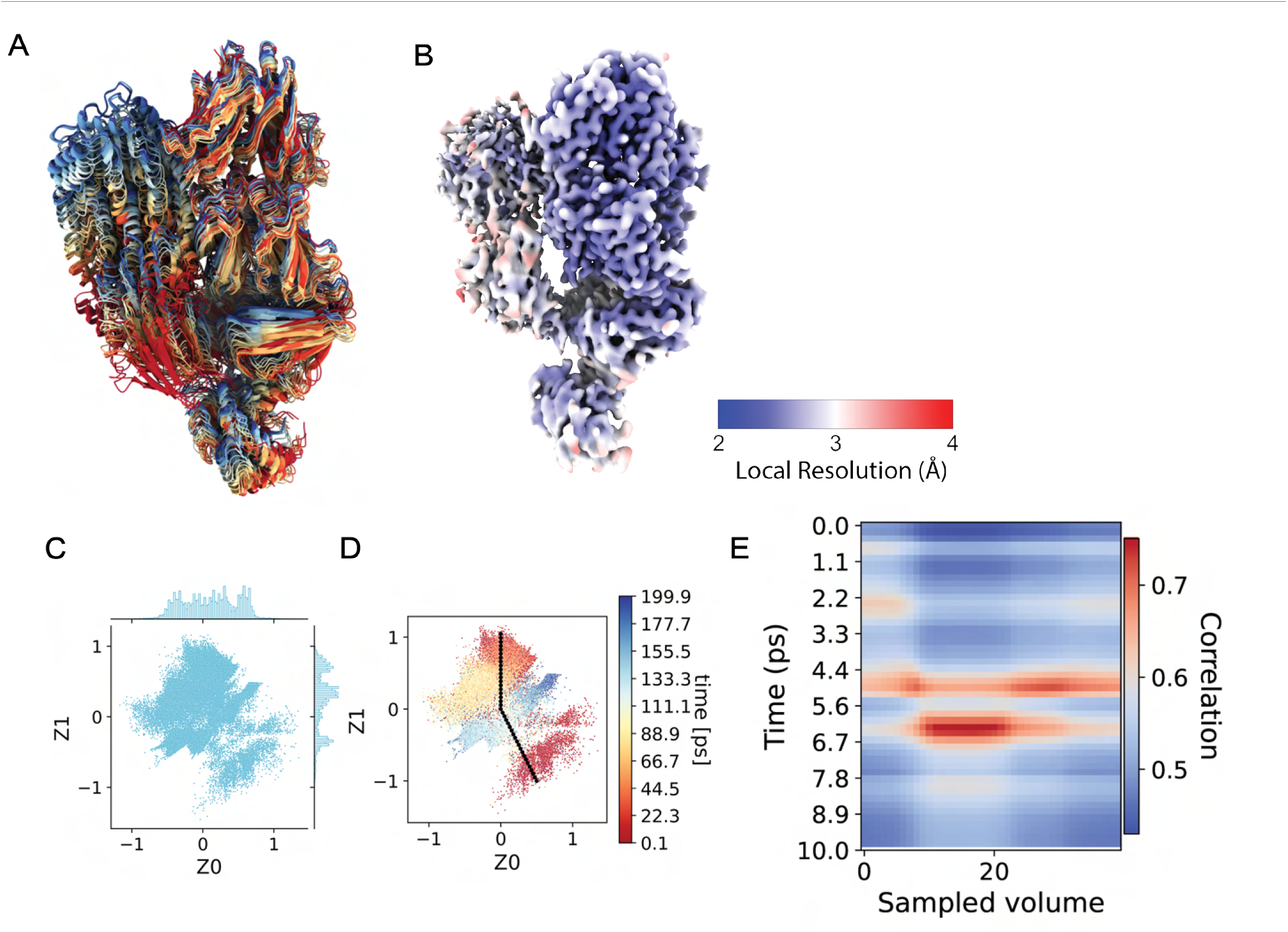
3DFlex training for C3-C3b steered MD with zdim = 2. **A**: Ensemble atomic model of the trajectory. Each model is coloured by the time in the MD simulation it originated from. **B**: Reconstructed consensus density map, coloured by local resolution. **C**: 2-dimensional latent space obtained. **D**: Latent space coloured by timepoint each particle originated from in the MD trajectory. Traversal through the latent space used for sampling volumes is plotted in black. **E**: Real-space map-to-model correlation heatmap between 50 states sampled from the MD trajectory and 41 sampled volumes.

**Figure S8.**
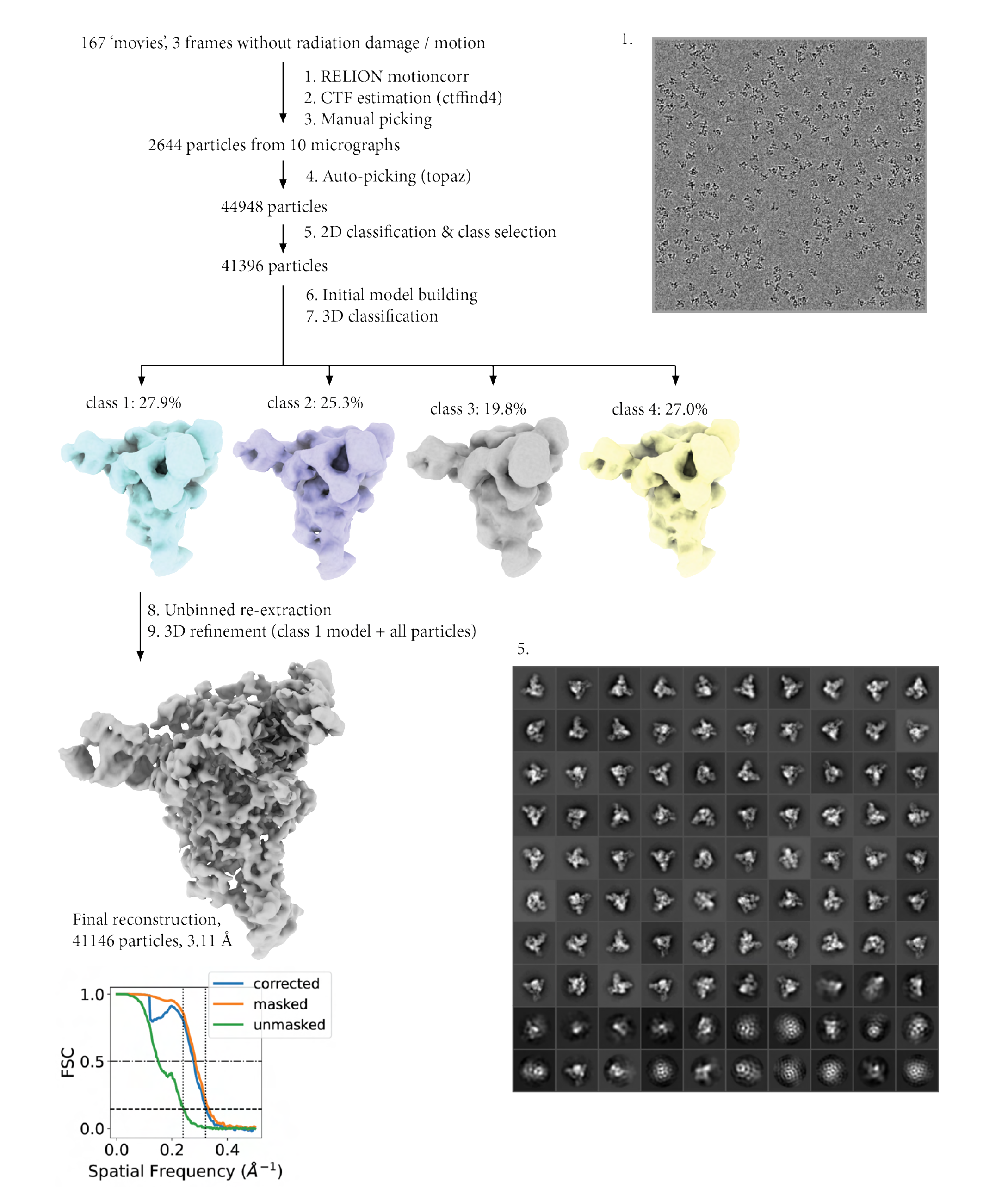
Processing workflow for SARS-CoV-2 RTC dataset based on DESRES-ANTON-13795965 MD trajectory.

**Figure S9.**
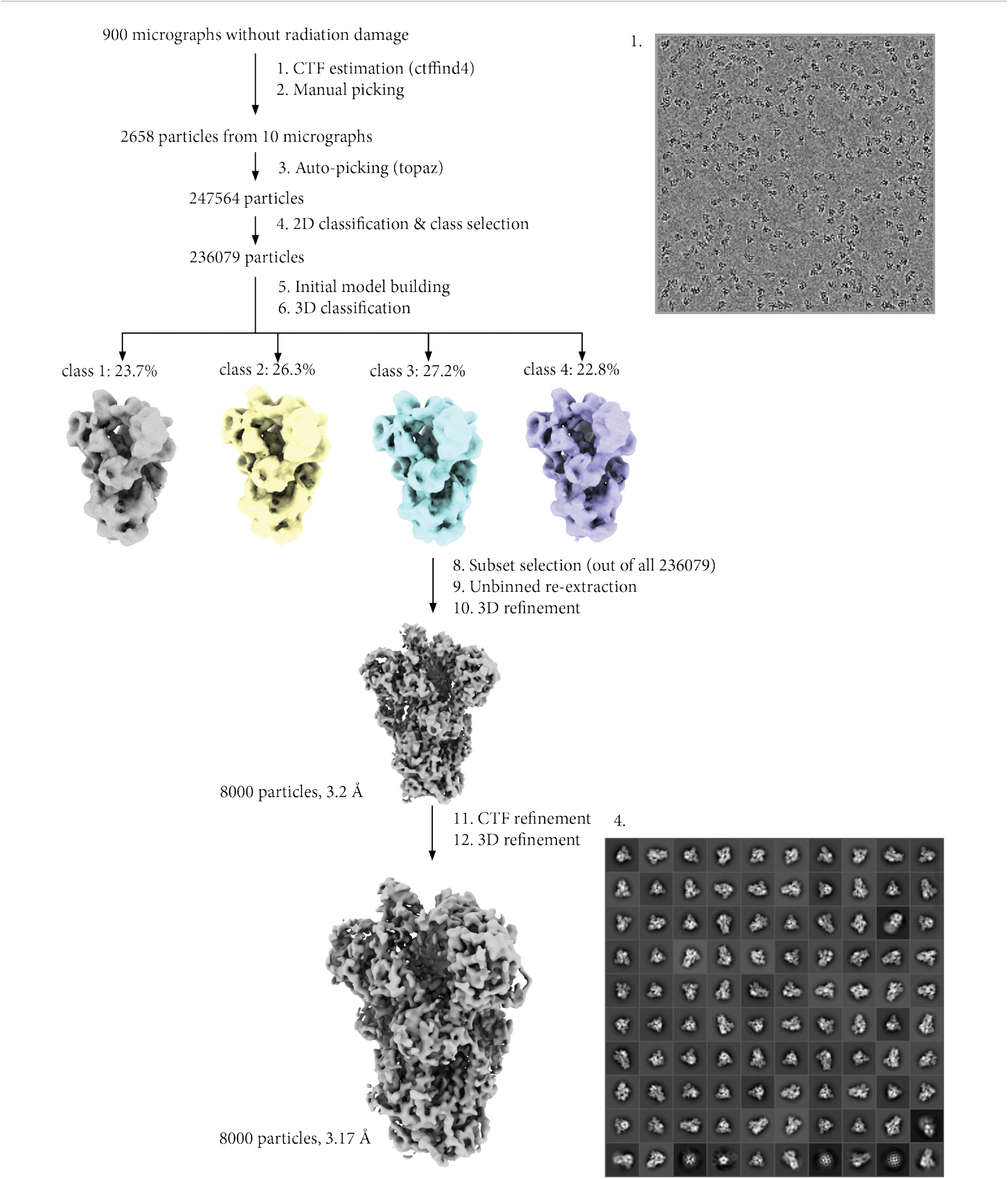
Processing workflow for SARS-CoV-2 spike glycoprotein dataset based on DESRES-ANTON-11021571 MD trajectory.

**Figure S10.**
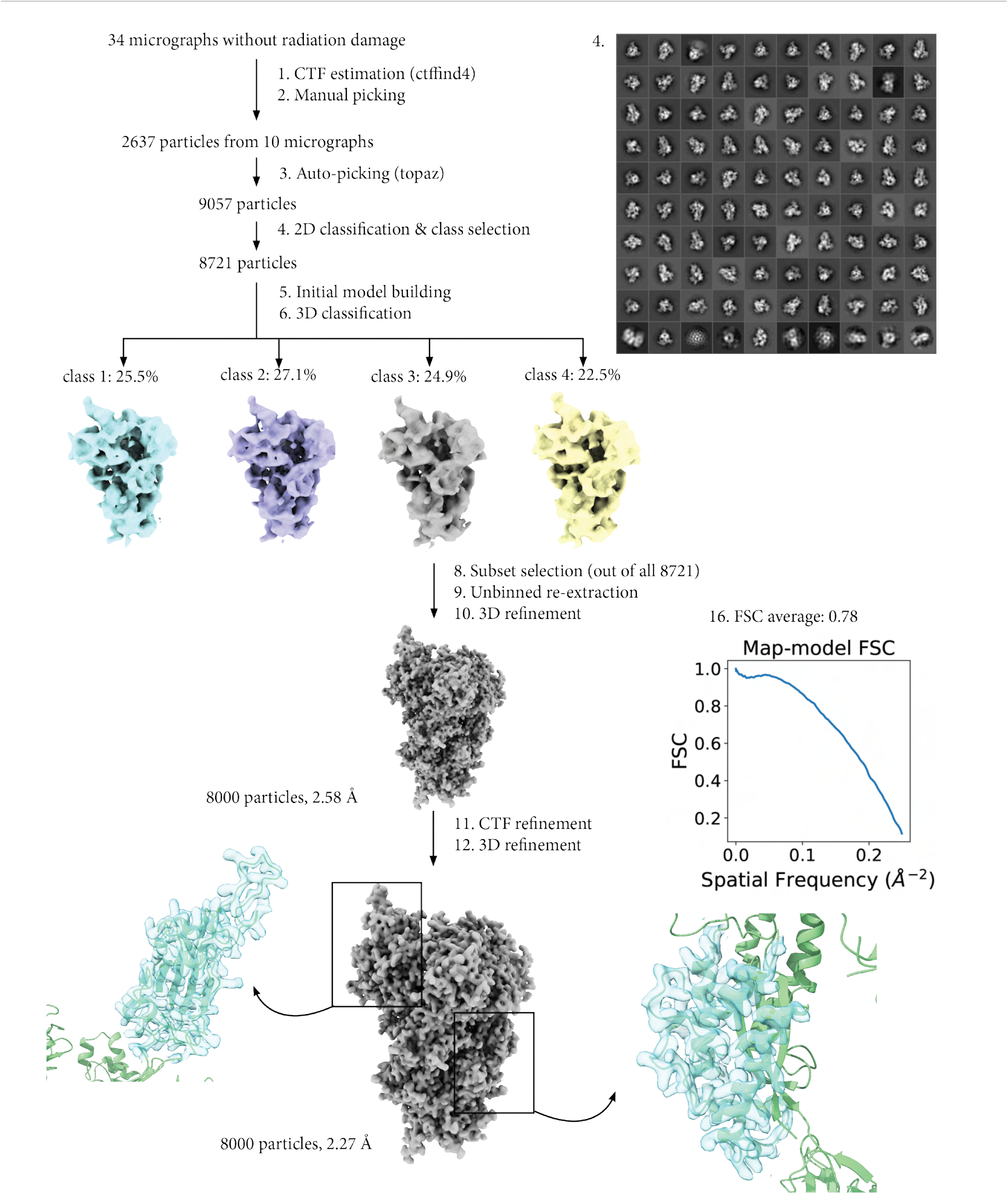
Processing workflow for SARS-CoV-2 spike glycoprotein dataset based on a single conformation from the DESRES-ANTON-11021571 MD trajectory.

**Figure S11.**
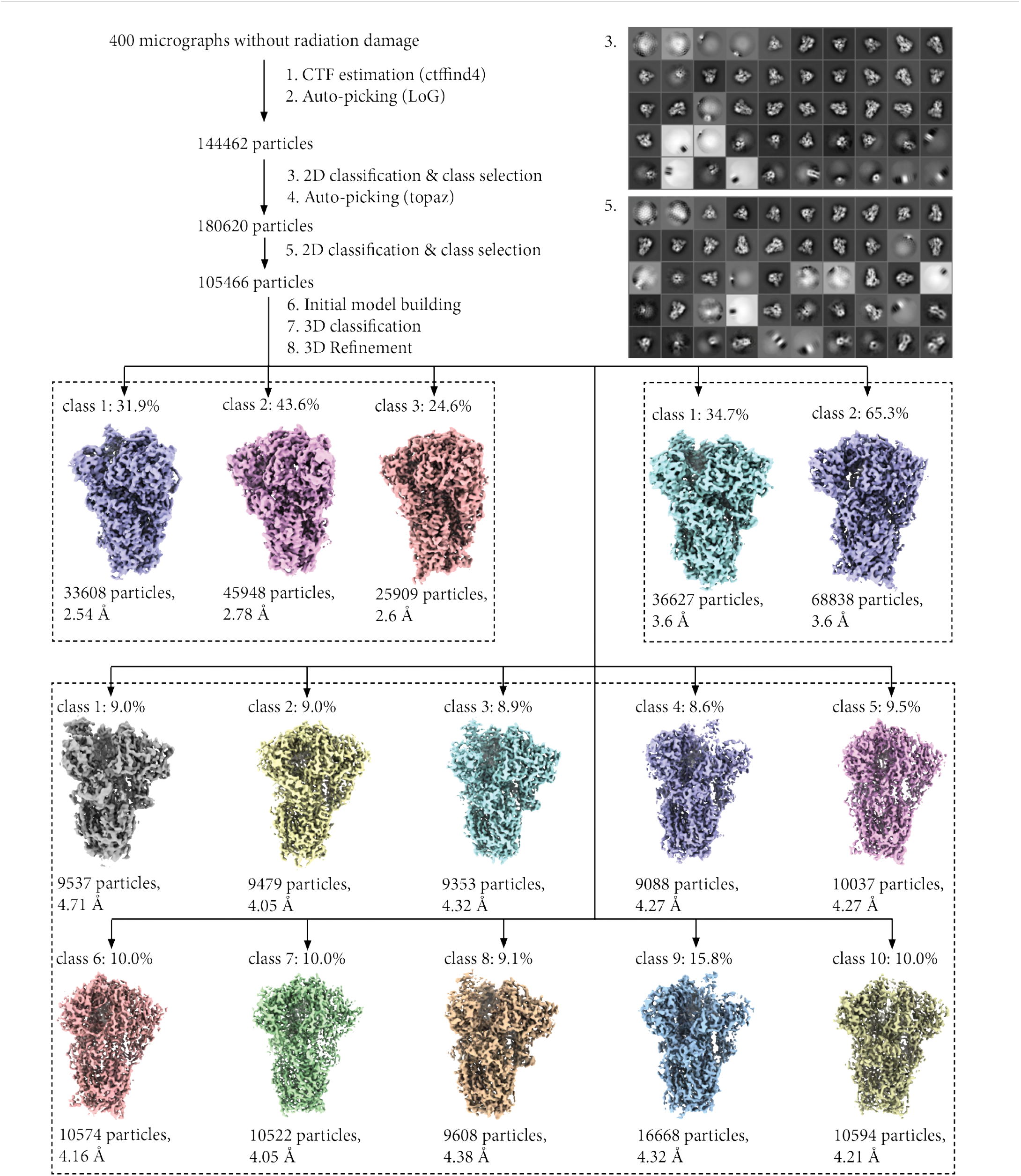
Processing workflow for mixed dataset based on the DESRES-ANTON-11021571 and DESRES-ANTON-11021566 trajectories. Density maps are shown for all classes after 3D classification with [2, 3, 10] classes and 3D refinement in RELION.

**Figure S12.**
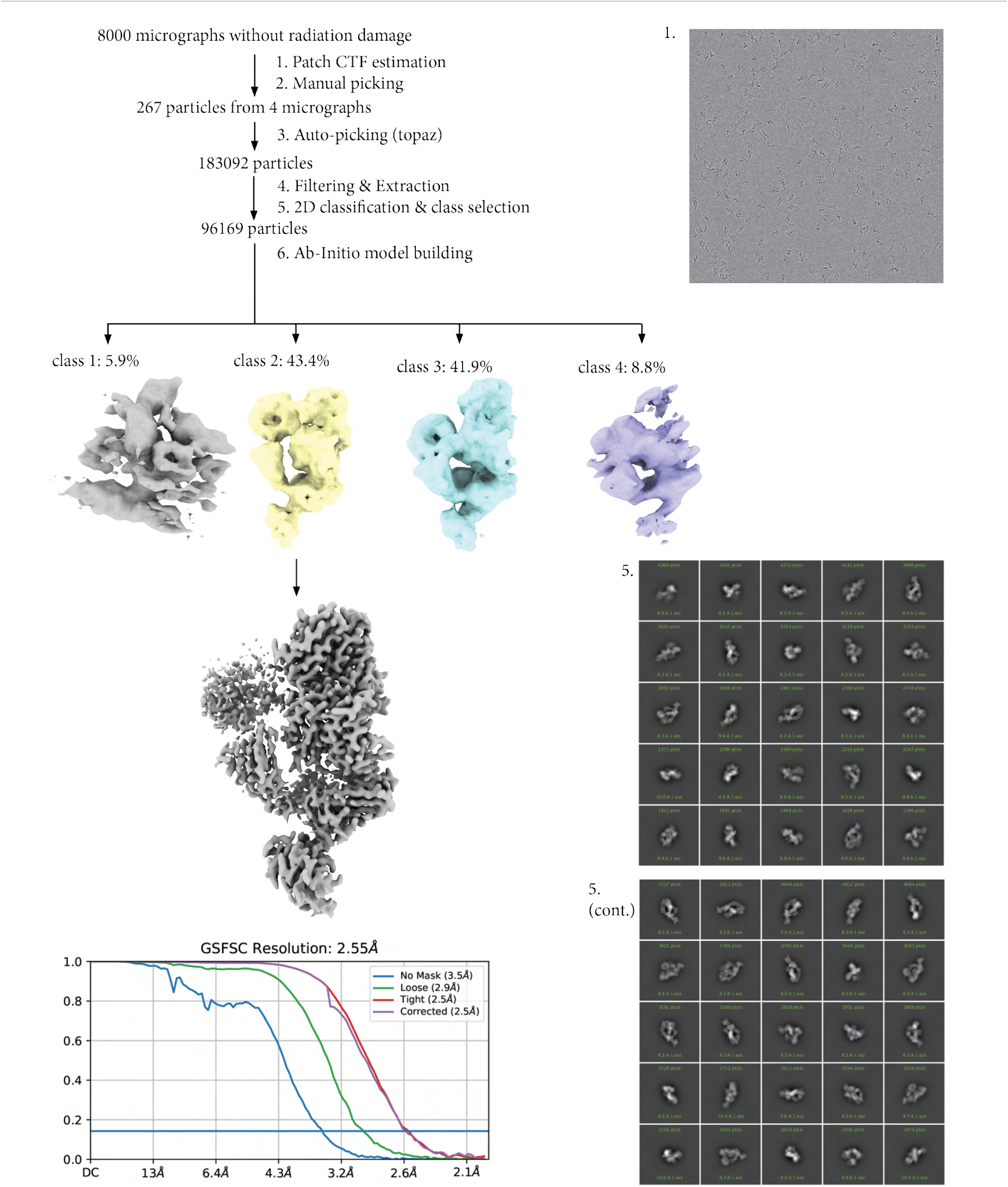
Processing workflow for C3 dataset based on steered MD simulation. Processing was done in CryoSPARC 4.2.1

**Figure S13.**
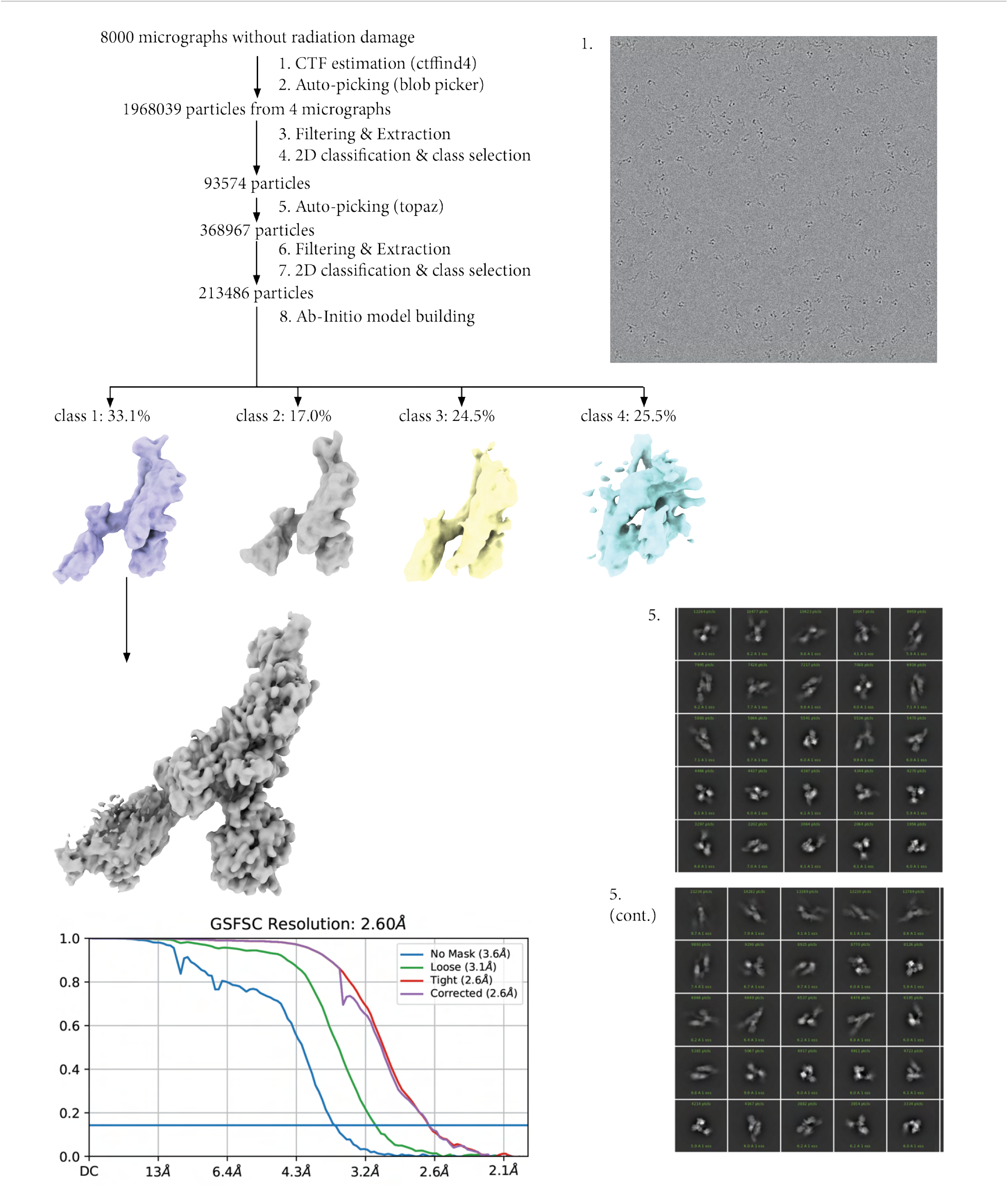
Processing workflow for single monomer SARS-CoV-2 spike glycoprotein based on steered MD simulation. Processing was done in CryoSPARC 4.2.1

## References

Beckers, M., Jakobi, A. J. & Sachse, C. (2019). IUCrJ, 6(1), 18–33.

Bepler, T., Morin, A., Rapp, M., Brasch, J., Shapiro, L., Noble, A. J. & Berger, B. (2019). Nature Methods, 16(11), 1153–1160. Publisher: Nature Publishing Group.

Bock, L. V. & Grubmüller, H. (2022). Nature Communications, 13(1), 1709. Number: 1 Publisher: Nature Publishing Group.

Chen, M. & Ludtke, S. J. (2021). Nature Methods, 18(8), 930–936. Number: 8 Publisher: Nature Publishing Group.

Cheng, Y. (2015). Cell, 161(3), 450–457.

Eastman, P., Swails, J., Chodera, J. D., McGibbon, R. T., Zhao, Y., Beauchamp, K. A., Wang, L.-P., Simmonett, A. C., Harrigan, M. P., Stern, C. D., Wiewiora, R. P., Brooks, B. R. & Pande, V. S. (2017). PLOS Computational Biology, 13(7), e1005659. Publisher: Public Library of Science.

Egelman, E. H. (2016). Biophysical Journal, 110(5), 1008–1012.

Frank, J. & Ourmazd, A. (2016). Methods, 100, 61–67.

Grant, T. & Grigorieff, N. (2015). eLife, 4, e06980. Publisher: eLife Sciences Publications, Ltd.

Greer, J., Joosten, M., Jakobi, A. & Burnley, T. (2024). ccpem/roodmus: v0.0.32. 10.5281/zenodo.10258256

Hayward, S. B. & Glaeser, R. M. (1979). Ultramicroscopy, 4(2), 201–210.

Huang, X., Wang, Y., Yu, C., Zhang, H., Ru, Q., Li, X., Song, K., Zhou, M. & Zhu, P. (2022). Science China Life Sciences, 65(12), 2491–2504.

Janssen, B. J. C., Christodoulidou, A., McCarthy, A., Lambris, J. D. & Gros, P. (2006). Nature, 444(7116), 213–216. Number: 7116 Publisher: Nature Publishing Group.

Kimanius, D., Dong, L., Sharov, G., Nakane, T. & Scheres, S. H. W. (2021). Biochemical Journal, 478(24), 4169–4185.

Kühlbrandt, W. (2014). Science, 343(6178), 1443–1444. Publisher: American Association for the Advancement of Science.

Leesch, F., Lorenzo-Orts, L., Pribitzer, C., Grishkovskaya, I., Roehsner, J., Chugunova, A., Matzinger, M., Roitinger, E., Belačić, K., Kandolf, S., Lin, T.-Y., Mechtler, K., Meinhart, A., Haselbach, D. & Pauli, A. (2023). Nature, 613(7945), 712–720. Number: 7945 Publisher: Nature Publishing Group.

Nakane, T., Kimanius, D., Lindahl, E. & Scheres, S. H. (2018). eLife, 7, e36861.

Nguyen, T. H. D., Galej, W. P., Bai, X.-c., Oubridge, C., Newman, A. J., Scheres, S. H. W. & Nagai, K. (2016). Nature, 530(7590), 298–302. Number: 7590 Publisher: Nature Publishing Group.

Noble, A. J., Wei, H., Dandey, V. P., Zhang, Z., Tan, Y. Z., Potter, C. S. & Carragher, B. (2018). Nature Methods, 15, 793–795.

Nogales, E. (2016). Nature Methods, 13(1), 24–27. Number: 1 Publisher: Nature Publishing Group.

Parkhurst, J. M., Cavalleri, A., Dumoux, M., Basham, M., Clare, D., Siebert, C. A., Evans, G., Naismith, J. H., Kirkland, A. & Essex, J. W. (2024). Ultramicroscopy, 256, 113882.

Parkhurst, J. M., Dumoux, M., Basham, M., Clare, D., Siebert, C. A., Varslot, T., Kirkland, A., Naismith, J. H. & Evans, G. (2021). Open Biology, 11(10), 210160.

Pintilie, G., Zhang, K., Su, Z., Li, S., Schmid, M. F. & Chiu, W. (2020). Nature Methods, 17(3), 328–334. Publisher: Nature Publishing Group.

Punjani, A. & Fleet, D. J. (2021). Journal of Structural Biology, 213(2), 107702.

Punjani, A. & Fleet, D. J. (2023). Nature Methods, 20(6), 860–870. Number: 6 Publisher: Nature Publishing Group.

Rohou, A. & Grigorieff, N. (2015). Journal of Structural Biology, 192(2), 216–221.

Rosenthal, P. B. & Henderson, R. (2003). Journal of Molecular Biology, 333(4), 721–745.

Scheres, S. H. W. (2016). In Methods in Enzymology, edited by R. A. Crowther, vol. 579 of The Resolution Revolution: Recent Advances In cryoEM, pp. 125–157. Academic Press. https://www.sciencedirect.com/science/article/pii/S0076687916300301

Schoppe, J., Schubert, E., Apelbaum, A., Yavavli, E., Birkholz, O., Stephanowitz, H., Han, Y., Perz, A., Hofnagel, O., Liu, F., Piehler, J., Raunser, S. & Ungermann, C. (2021). Journal of Biological Chemistry, 297(5), 101334.

Schwab, J., Kimanius, D., Burt, A., Dendooven, T. & Scheres, S. H. W. (2023). DynaMight: estimating molecular motions with improved reconstruction from cryo-EM images. https://www.biorxiv.org/content/10.1101/2023.10.18.562877v1

Serna, M., González-Corpas, A., Cabezudo, S., López-Perrote, A., Degliesposti, G., Zarzuela, E., Skehel, J., Muñoz, J. & Llorca, O. (2022). Nucleic Acids Research, 50(2), 1128–1146.

Shaw, D. E. (2020). Molecular Dynamics Simulations Related to SARS-CoV-2. https://www.deshawresearch.com/downloads/downloadtrajectorysarscov2.cgi/

Stagg, S. M., Noble, A. J., Spilman, M. & Chapman, M. S. (2014). Journal of Structural Biology, 185(3), 418–426.

Yang, Y. I., Shao, Q., Zhang, J., Yang, L. & Gao, Y. Q. (2019). The Journal of Chemical Physics, 151(7), 070902.

Zhong, E. D. (2022). Machine Learning for Reconstructing Dynamic Protein Structures from Cryo-EM Images. Phd thesis, Massachusetts Institute of Technology.

Zhong, E. D., Bepler, T., Berger, B. & Davis, J. H. (2021). Nature Methods, 18(2), 176–185. Number: 2 Publisher: Nature Publishing Group.

Zhou, T., Tsybovsky, Y., Gorman, J., Rapp, M., Cerutti, G., Chuang, G.-Y., Katsamba, P. S., Sampson, J. M., Schön, A., Bimela, J., Boyington, J. C., Nazzari, A., Olia, A. S., Shi, W., Sastry, M., Stephens, T., Stuckey, J., Teng, I.-T., Wang, P., Wang, S., Zhang, B., Friesner, R. A., Ho, D. D., Mascola, J. R., Shapiro, L. & Kwong, P. D. (2020). Cell Host & Microbe, 28(6), 867–879.e5.

